# The Pursuit of Pride: Outcomes achieved under Beliefs of Internal Control shape positive Affect and neural Dynamics in the vmPFC

**DOI:** 10.1101/637207

**Authors:** David S Stolz, Laura Müller-Pinzler, Sören Krach, Frieder M Paulus

## Abstract

Experiencing events as controllable and attributing positive outcomes to own contributions is essential for human well-being. Based on classic psychological theory we test how internal control beliefs impact the affective valuation of task outcomes, neural dynamics and ensuing behavioral preferences. In three consecutive studies with independent samples we show that dynamics in self-evaluative affect specifically increase when agents believe they caused a given task outcome. We demonstrate that these outcomes engage brain networks processing self-referential information in the cortical midline. Here, activity in the ventromedial prefrontal cortex tracks outcome valence regarding both success as well as internal control, and covaries with self-evaluative affect. These affective dynamics also relate to increased functional coupling between the ventral striatum and cortical midline structures. Finally, we show that self-evaluative affect promotes preferences for control, even at a monetary cost. Our investigations extend recent models of positive affect and well-being, and emphasize that control beliefs drive intrinsic motivation.

## Introduction

The subjective belief of being in control over events in one’s life is essential for well-being^1–3^. The ways in which individuals perceive themselves or external forces as determining their fate has been conceptualized in the classic theory of locus of control^4^ and since then has deeply influenced theory formation in psychology^5–7^. Here, the sense of being in control hinges on the subjective belief that the course of events can be shaped by own efforts and actions (i.e. internal control beliefs), creating the cognitive foundation of attributing their outcomes to the self. Internal control beliefs can determine whether individuals will show effort^8^, make career choices^9^, and are also more generally considered to be a protective factor for various psychiatric phenomena^10–12^. In contrast, the conviction of having no control and thus being at the mercy of chance or other forces in one’s environment (i.e. external control beliefs) has been linked to learned helplessness and major depression^13^, a debilitating psychiatric condition that is characterized by attenuated affect, reduced levels of motivated behavior and diminished self-esteem^14^. In respect of this background, we aimed to investigate how contexts that foster the formation of internal control beliefs and self-attribution shape the affective valuation, neural processing, and motivational consequences of task outcomes.

Having opportunities to choose is a fundamental prerequisite of exerting control^3^. Choice opportunities allow to pick between options with different value^15^, and thus are a means for maximizing the probability of achieving desired outcomes, avoiding potential hazards, and reducing uncertainty^16^. The importance of exerting control over one’s environment has stimulated the idea that having choices per se carries intrinsic value^3^. Studies have demonstrated that humans favor choice options that are followed by a second choice over those that are not^17^, or prefer tasks with more over those with fewer choice options^18^. Relatedly, humans prefer stimuli selected freely during a preceding task over those that a computer had selected for them, despite equal monetary value of the stimuli^19^. Prior neuroscience studies also support the notion that choices have intrinsic value. Precisely, cues signaling an upcoming choice are associated with increased activity in the ventral striatum (VS)^20, 21^, a region implicated in dopaminergic reward processing^22–24^.

Although having choices is the condition precedent for exerting control, the lynchpin of theory and empirical evidence relating control beliefs to well-being are self-related thoughts and subjective models of whether an outcome can be achieved due to the capabilities of the agent^1, 2, 4, 6^. Thereby, the experience of control should increase if internal models of behavior are available that explain how actions – if executed correctly – yield a specific outcome^25, 26^, in contrast to settings in which behavior is thought to be linked to its outcomes by chance alone^20, 21^. One could consider skilled players in the game of darts who perform at a success rate of exactly 50% for hitting the bullseye. According to theory^4^, these players should to some extent experience internal control in a sense that they know they can shape the outcomes of the game, as their odds for hitting their target by far exceed those of a random thrower or less skilled player. Here, internal control beliefs set the cognitive foundation for being able to attribute the outcomes to own contributions and capabilities. However, when placing a bet on a color in a game of roulette, control beliefs should diminish as behavior and outcomes are linked only by chance even though outcome probabilities are similar to the previous example. Thus, while having choices is the condition necessary for exerting control, they are in principle not sufficient for experiencing control as implied by theory^4, 6, 7^. Only if the context offers internal control over outcomes, it is possible to attribute events to the self, own efforts, and abilities, with broad implications for self-related affect and motivation^27, 28^.

Theories on self-conscious affect predict that control beliefs shift the valuation of outcomes and result in distinct patterns of affective experience^29^. Precisely, the attribution of outcomes to controllable and internal causes is a necessary condition for the experience of the self-conscious affect of pride^27^. The concept of pride essentially hinges on a subjective model of control – the belief that an outcome was caused by one’s own actions – resulting in self-approval in case the event is relevant for personal goals^30, 31^. Pride experiences thus underlie self-esteem^32^, foster intrinsic motivation^28^, and mediate the contribution of internal control beliefs to well-being^2, 32^. In contrast, positive events caused by uncontrollable factors such as winning a lottery also drive affective dynamics such as momentary happiness^33, 34^, but do not have similar consequences in terms of self-relevance and motivation^28^. In this line, earnings that have been obtained through own efforts and labor are valued more than windfall gains^35^, thereby contributing to the effort paradox that the value of effort sometimes exceeds its costs^36^.

In the present study we test how the affective valuation of outcomes varies according to how much a task allows for developing internal control beliefs and self-attribution, how this relates to shifts in neural processing of objectively similar task outcomes and we finally provide evidence that the subjective value of tasks with a high degree of control is related to the dynamics of self-evaluative affect. To do so, we introduce a novel paradigm that in addition to mere choice^20^ extends the concept of control to an experimental condition allowing subjects to attribute task outcomes to their own performance^26^, while strictly controlling outcome probabilities. This is a key step since with increasing control beliefs, outcomes also get more predictable and less uncertain in everyday life^6, 7, 37, 38^.

As predicted by theory, affective and especially pride dynamics should be most strongly modulated by outcomes if these are perceived to depend on one’s own behavior. Aside from the ventral striatum, prior evidence suggests the ventromedial prefrontal cortex (vmPFC) to be a key cortical interface for integrating self-attribution during outcome valuation. The vmPFC has extensive connections with the ventral tegmental area, the amygdala and the striatum^39, 40^, is associated with domain general computations of value^15, 41^ and has been linked to behavioral control and persistence in the face of failure^42–44^. Further, the vmPFC belongs to structures of the cortical midline implicated in self-related processing (CMS)^45, 46^. Specifically, the vmPFC has been associated with the generation of affective meaning^47, 48^, the value of revealing information about the self^49^, and updating of self-esteem and self-relevant optimistic beliefs^50, 51^. These findings suggest the vmPFC to serve a central function for the self-relevance of task outcomes under varying levels of control and associated self-related affective dynamics.

In order to manipulate internal control beliefs and investigate its impact on affective, neural, and motivational dynamics, we present a series of three consecutive experiments with independent samples (total *N*=129). In a novel paradigm we systematically manipulate control beliefs beyond having a choice or not, inducing low, medium, and high levels of perceived control (LC, MC, HC) via, respectively, a non-choice task, a choice task, and a skill-based task that allows attributing outcomes to the self. In principle, the choice and the non-choice task are gambles, while the skill-based task suggests participants to outperform chance level and having the capabilities needed to discriminate basic stimulus properties. This should imply that participants develop different degrees of control beliefs across conditions. Across all three conditions, rates of successes (WIN) and non-successes (noWIN) are kept at 50% (see figure 1). In study 1 we test whether affective responses to outcomes depend on internal control beliefs as predicted by theory. In study 2, we replicate and extend study 1 by characterizing affective and neural dynamics in response to outcomes obtained at different levels of internal control during task execution. We focus our analyses on brain regions processing reward value and self-referential processes, specifically the vmPFC. As a last step, in study 3 we predict the subjective value controllable environments based on the dynamics of self-related positive affect by introducing an adaptive preference task. Our results support the broad impact of internal control beliefs on affect, motivated behavior and underlying brain function. We find that the value of control is associated with self-related affective responses and may even outweigh a monetary detriment entailed by preferences for control.

**Figure 1.**
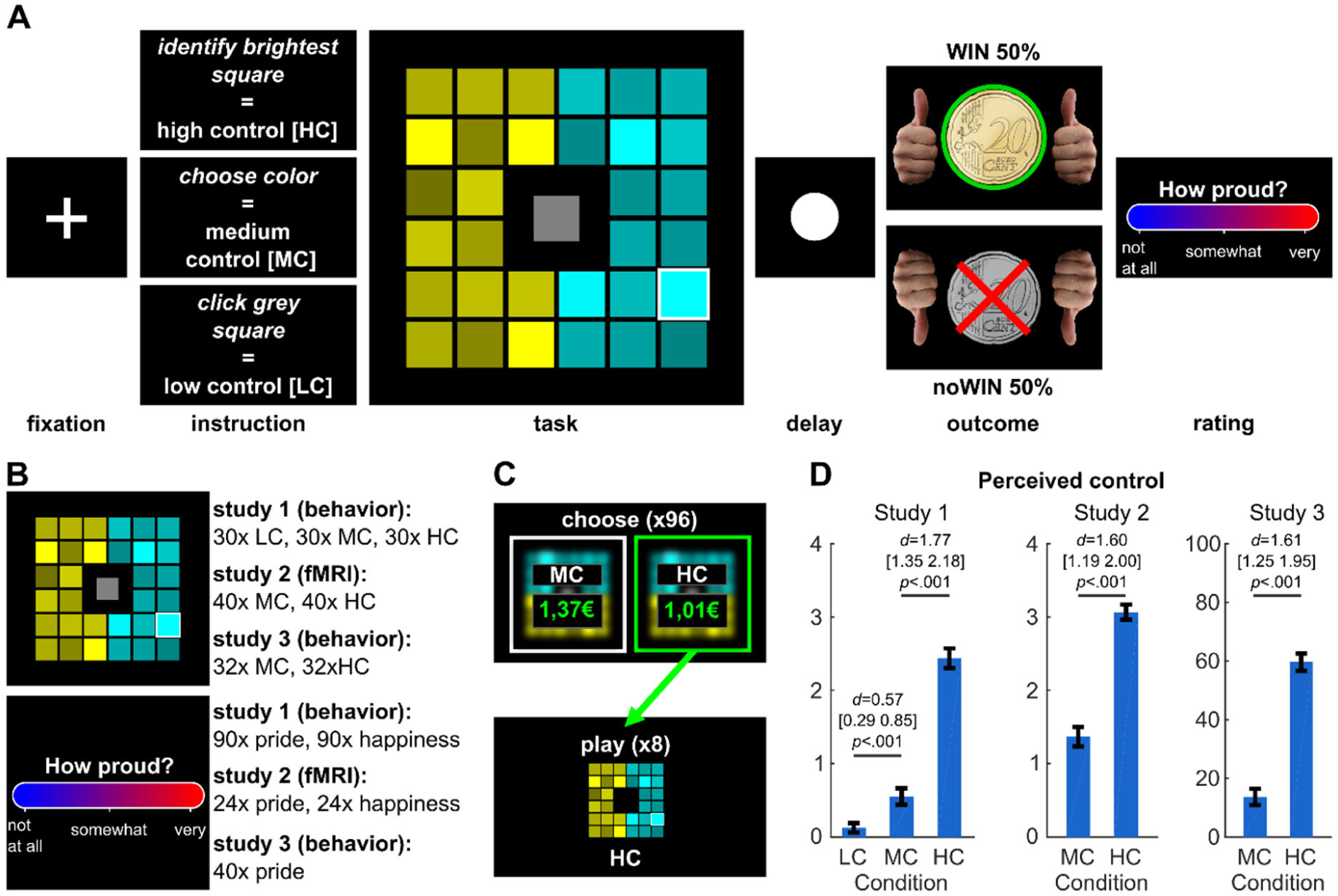
Overview about the experimental paradigm and the three studies. **A** Trial structure of the task. A fixation cross was followed by a cue indicating that the participant has to find the brightest square within either turquoise or yellow (high control, HC), choose between yellow and turquoise (medium control, MC), or play a gamble by clicking on the grey square in the middle (low control, LC; only presented in study 1). After the cue, the task grid occurred and was followed by a delay and task outcomes. Pride and happiness ratings regarding the preceding outcome were presented after the outcome phase. **B** Overview of the three studies. The task was presented in each study with varying numbers of trials and conditions. **C** In study 3, after the main task, 96 trials of a choice task were presented. The choice options offered to play either HC or MC with adaptively changing amounts of money determined using a staircase algorithm. Participants were informed that some of the chosen options would be actually presented and that the associated monetary gain would be awarded at the end of the experiment. **D** Internal control belief ratings for each study (‘How much could you influence the outcome of the task’). *d* = Cohen’s *d*. Values in brackets represent lower and upper bounds of 90% confidence intervals. HC=high control. MC=medium control. LC=low control. Error bars are +/− 1 standard error of the mean.

## Results

### Study 1: Dynamics of Self-Evaluative Positive Affect

Study 1 manipulated levels of internal control beliefs above and beyond having a choice or not. In brief, we presented participants (*N*=40) with a grid of one centered grey square and 32 surrounding squares that were separated into two parts of shaded colors (see figure 1). This allowed to implement three different conditions that were pseudorandomly presented and cued before each trial. On the lowest level of control, successes (WIN outcomes; 0.20 €) and non-successes (noWIN outcomes; 0.00 €) depended on an automated gamble which was initiated by clicking on the grey square (low control, LC; see figure 1). On the medium level of control beliefs, WIN and noWIN outcomes allegedly depended on the correct choice between the two colors (medium control, MC). On the highest level of internal control, WIN and noWIN outcomes depended on whether subjects were able to identify the brightest square within the shades of one color (high control, HC). Unknown to the participants, across all three conditions outcome probabilities were held constant at 50% by predefined feedback and participant learned that their hit rate was 50% on average in the HC condition (see methods for details). Each condition was presented 30 times, i.e. a total of ninety trials was presented, and ratings of both pride and happiness were acquired after each outcome. After completion of the experimental tasks, subjects rated their control beliefs together with subjective experience of outcome probabilities for the different conditions.

Subjects experienced the HC condition as more controllable than the other conditions (main effect of condition: *F*(1.74, 76.72)=133.97, *p*<.001; planned comparisons: HC>MC: *t*(39)=11.21, *p*<.001, *d*=1.77, 90% CI=[1.35; 2.18]; MC>LC: *t*(39)=3.60, *p*<.001 *d*=0.57, 90% CI=[0.29; 0.85]; Bonferroni-corrected for multiple comparisons, one-sided). As predicted by theory, the dynamics in the pride experiences induced by succeeding (WIN vs noWIN) depended more strongly on the task than it was the case for happiness ratings (condition*affect*outcome valence: *F*(1.29,50.10)=11.44, *p*<.001, 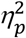=.23; figure 2, for full rmANOVA effects, see Supplement 1 – table S1). Paired comparisons showed that affect ratings were higher for WIN than for noWIN outcomes for both happiness (LC: *t*(39)=7.85, *p*<.001; MC: *t*(39)=8.16, *p*<.001; HC: *t*(39)=9.74, *p*<.001) and pride (LC: *t*(39)=3.29, *p*<.001; MC: *t*(39)=4.41, *p*<.001; HC: *t*(39)=7.37, *p*<.001, all *p*-values one-sided and Bonferroni-corrected for multiple comparisons). These differences were larger in HC than in MC (pride: *t*(39)=6.97, one-sided *p*<.001; happiness: *t*(39)=4.64, two-sided *p*<.001), and larger in MC than in LC for pride (*t*(39)=3.03, one-sided *p*=.013), but not happiness (*t*(39)=2.24, two-sided *p*=.093, corrected), showing that internal control beliefs drive affective dynamics in general. Importantly, the pride response (i.e. the difference [HC:WIN-HC:noWIN]-[MC:WIN-MC:noWIN]) increased more strongly from MC to HC than the happiness response (*t*(39)=3.69, one-sided *p*<.001), while no difference was found when comparing pride and happiness responses between LC and MC (*t*(39)=1.57, one-sided *p*=.124, corrected). These findings demonstrate that control beliefs depend on contexts in which behavior can be guided by subjective models that explain how outcomes can be achieved through own actions and the observed performance can only be achieved by a certain degree of mastering the task. If such self-attribution is possible, internal control increases beyond having a choice or not, effectively increasing the self-relevance of outcomes and changing their affective valuation, driving pride more strongly than happiness^27^. In contrast, merely having a choice may be rewarding in itself^20^, but falls short of internal control beliefs as defined in classic theories and has little impact on self-evaluative affect.

**Figure 2.**
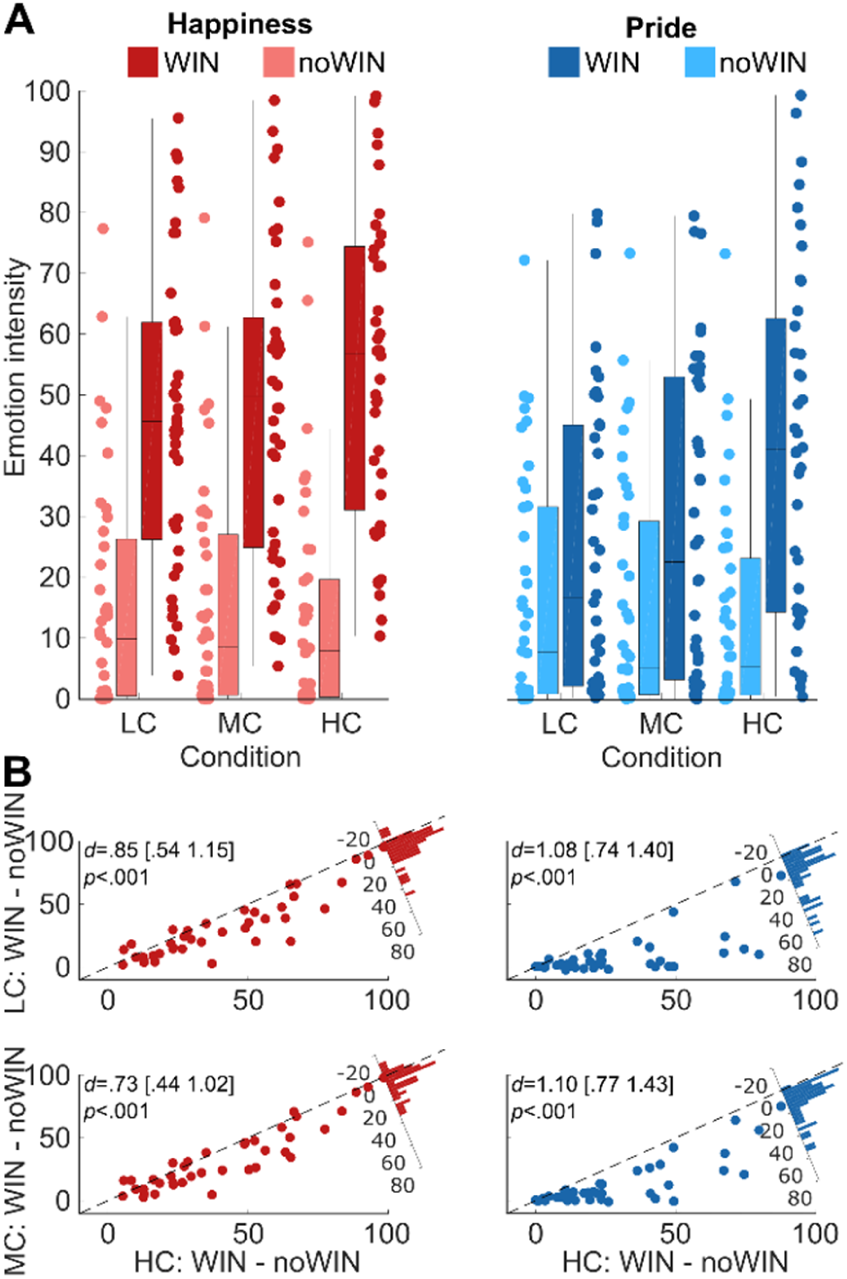
Task effects on emotion ratings in study 1 **A** Emotion ratings from study 1, separate for pride and happiness, high, medium and low control as well as outcome valence. **B** Scatterplots depict reactivity to WIN vs noWIN outcomes for happiness (left) and pride (right) in dependence of perceived control (x-axis: HC; y-axis (top): LC; y-axis (bottom): MC). Top and bottom rows depict relationships between emotion reactivity in LC and HC, as well as MC and HC, respectively. Histograms in top right corners visualize the differences in emotion reactivity between HC and LC (top row), or HC and MC (bottom row). HC=high control. MC=medium control. LC=low control. *d*=Cohen’s *d*. Values in brackets represent 90% confidence intervals for *d*.

### Study 2: Neural Foundations of Outcome Valuation in the Context of Internal Control

In study 2 we replicated and extended study 1 by characterizing the neural processing of outcomes achieved under internal control and their impact on affective dynamics. Subjects (*N*=39) underwent fMRI scanning while performing the identical MC and HC conditions from study 1, which there showed the strongest modulation of affect and are most relevant for developing theory and the neuroscience of control beliefs. A total of 80 trials were presented, with ratings of either pride or happiness (24 each) acquired after every second to third outcome.

#### Dynamics of self-evaluative positive Affect

The results of study 2 replicated those of study 1 (figure 3). Subjects experienced much higher internal control in HC than in MC (paired-sample t-tests: *t*(37)=9.85, *p*<.001, *d*=1.60, 90% *CI*=[1.19; 2.00]), and affect ratings differed between tasks with different levels of control (rmANOVA: main effect of condition: *F*(1,38)=8.38, *p*=.006, 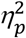=.18). Comparably to study 1, pride responses to task outcomes differed from those for happiness in dependence of control beliefs (condition*outcome valence*affect: *F*(1,38)=9.08, *p=*.005, 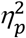=.19). As before, we computed the affective responses separately for pride and happiness ([HC:WIN–HC:noWIN] – [MC:WIN–MC:noWIN]). The resulting pride response was significantly larger than the happiness response (*t*(38)=3.01, one-sided *p*=.005), emphasizing how self-related positive affect hinges on the subjective belief that outcomes depend on one’s own actions (for full rmANOVA effects, see Supplement 1 – table S2).

**Figure 3.**
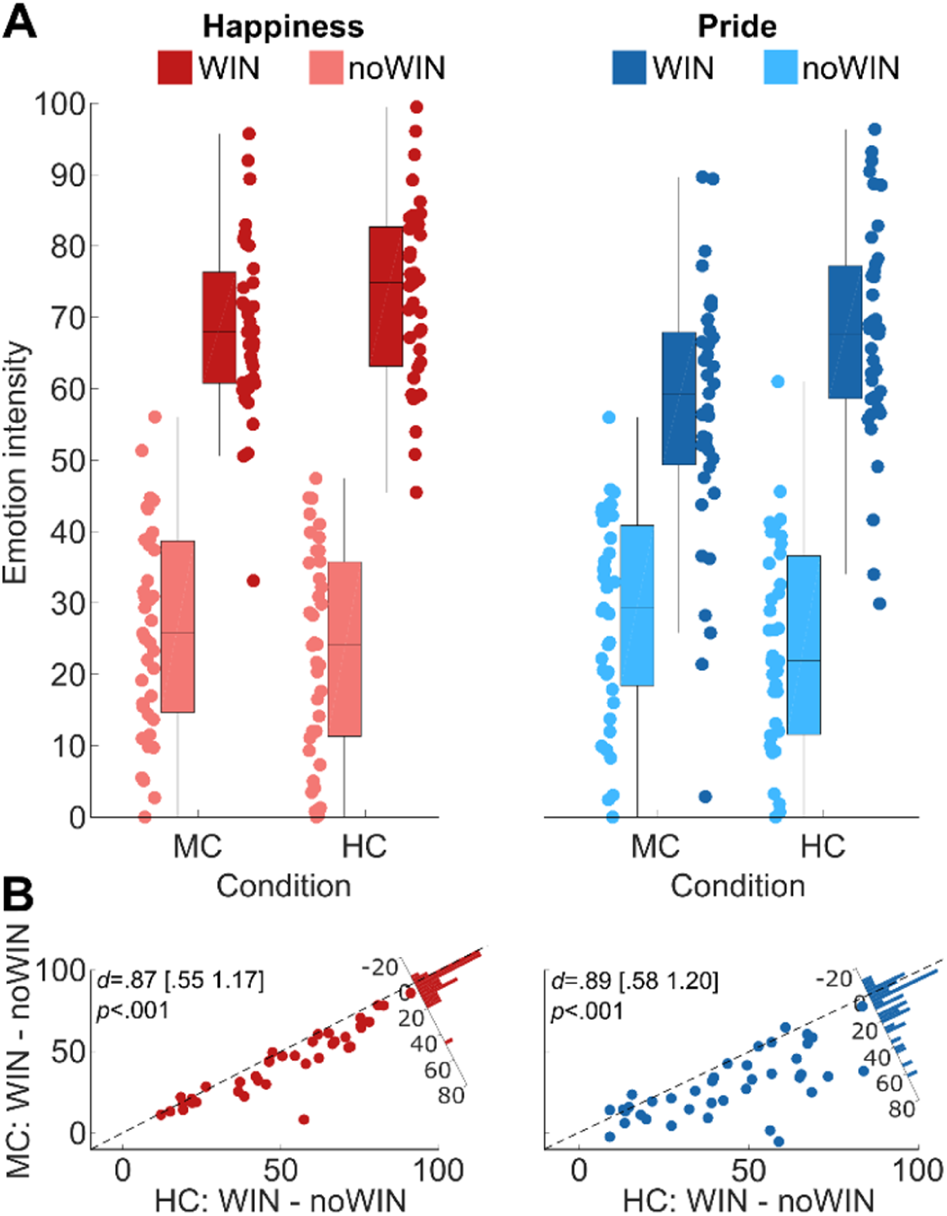
Task effects on emotion ratings in study 2 **A** Emotion ratings from study 2, separate for pride and happiness, high and medium control as well as outcome valence. **B** Scatterplots depict reactivity to WIN vs noWIN outcomes for happiness (left) and pride (right) in dependence of internal control condition (x-axis: HC; yaxis: MC). Histograms in top right corners visualize the differences in emotion reactivity between HC and MC. HC=high control. MC=medium control. *d*=Cohen’s *d*. Values in brackets represent 90% confidence intervals

**Figure 4.**
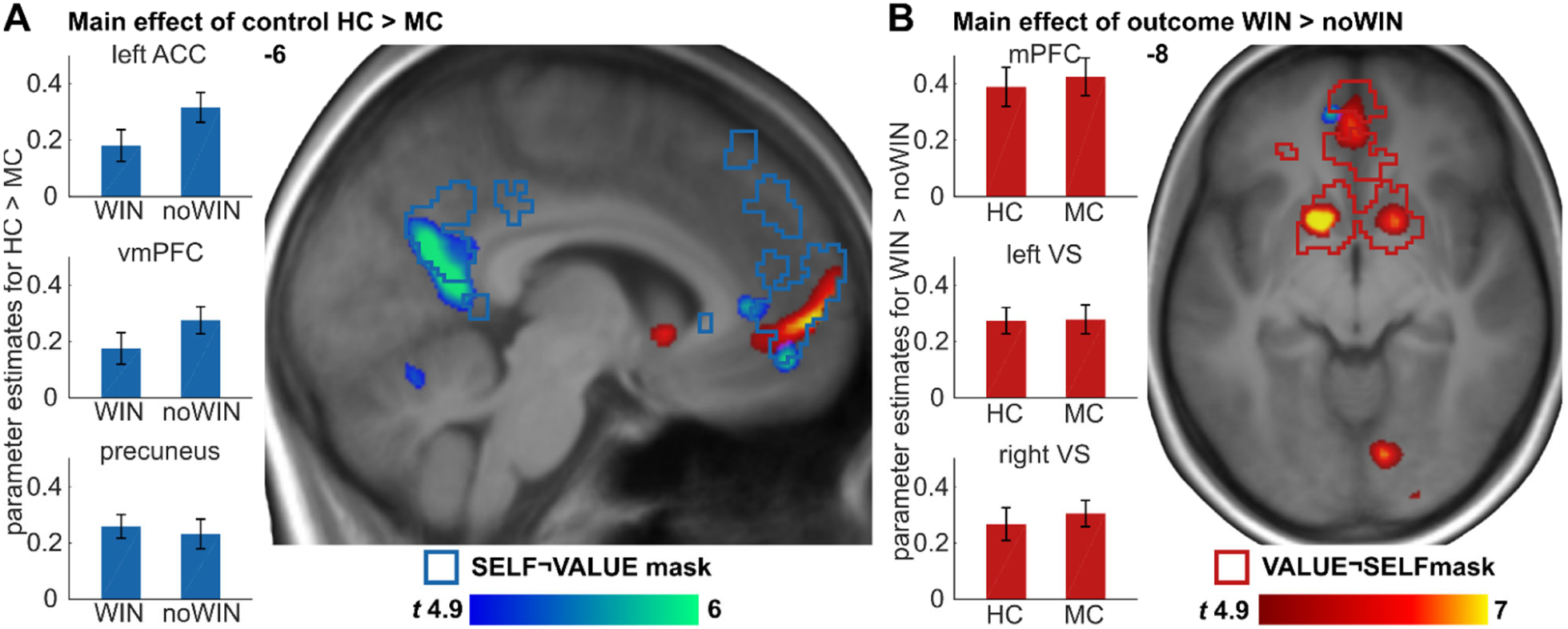
BOLD effects of the MRI task. **A** HC>MC (blue-green) and **B** WIN>noWIN (red-yellow) displayed at *p*<.05, whole-brain FWE-corrected. Blue and red outlines represent masks for SELF¬VALUE and VALUE¬SELF, respectively. Bar plots on the left represent average effects across participants for all voxels inside HC>MC clusters. Bar plots on the right depict average effects across participant s for all voxels inside WIN>noWIN clusters. Error bars are +/− 1 standard error of the mean.

#### Neural Responses to Outcomes in the Context of Internal Control

Receiving a positive outcome (WIN>noWIN) yielded significant activations in regions associated with processing of value (as indicated by ROI-analyses using the VALUE¬SELF mask, covering areas associated with the term “value”, but not “self-referential”; created using Neurosynth^52^; see methods). Specifically, we observed significant activation with peaks in bilateral ventral striatum (VS; left: x,y,z (mm): −14,8,−10, *t*(114)=8.18; *Z*=7.24, *k*=235; right: 16,4,−12, *t*(114)=7.69; *Z*=6.89, *k*=233; all coordinates in MNI space), anterior cingulate cortex (ACC; −2,44,−6, *t*(114)=6.71; *Z*=6.15, *k*=108), superior medial frontal gyrus (−2,58,0, *t*(114)=8.14; *Z*=7.21, *k*=61), and vmPFC (4,48,−2, *t*(114)=5.59; *Z*=5.24, *k*=12; all p-values <.05, FWE-corrected within the VALUE¬SELF mask; for whole-brain effects see Supplement 2).

Receiving outcomes attributable to one’s own actions (HC>MC) yielded significant activations in CMS (ROI-analysis using the SELF¬VALUE mask, covering regions associated with the term “self-referential” but not “value”), with peaks in left vmPFC (−8,50,−12, *t*(114)=6.34; *Z*=5.85, *k*=27), and left ACC (−2,38,4, *t*(114)=6.27; *Z*=5.80, *k*=108). In addition, this contrast yielded significant effects in posterior CMS, specifically the left cuneus (−8,−58,20, *t*(114)=6.61; *Z*=6.07, *k*=208), extending to precuneus (−6,−50,6, *t*(114)=5.60; *Z*=5.25, *k*=14; all *p*-values <.05, small-volume FWE-corrected). These findings indicate, that, as hypothesized, cortical regions associated with processing of self-relevant information are engaged when outcomes are obtained in task environments that are perceived as controllable and thus provide information about the subjects’ capabilities^31, 53^.

Next, we were interested in the additive effects of internal control and winning, by describing specific neural correlates of receiving positive outcomes under high control beliefs (three-way conjunction of [HC:WIN>HC:noWIN] ∩ [HC:WIN>MC:WIN] ∩ [HC:WIN>MC:noWIN]; figure 5). This analysis revealed that the left vmPFC, specifically, is more strongly engaged when subjects’ successes are attributable to their own actions (−4,48,−10, *t*(114)=3.45; *Z*=3.36, *k*=11, *p*=.023, FWE-corrected within the SELF∩VALUE mask, covering regions associated with both “self-referential” and “value”). The additivity of the effect in this region is illustrated by the common of responses to winning (WIN>noWIN: −2,48,−4, t(114)=7.60; Z=6.82, k=193; p<.05, FWE-corrected) as well as task controllability (HC>MC; −8,48,−12, t(114)=6.13; Z=5.69, k=162, p<.05, small-volume FWE-corrected). The localization of this effect thus mirrors how the processing of value and self-reference converges in the vmPFC^51, 54^.

**Figure 5.**
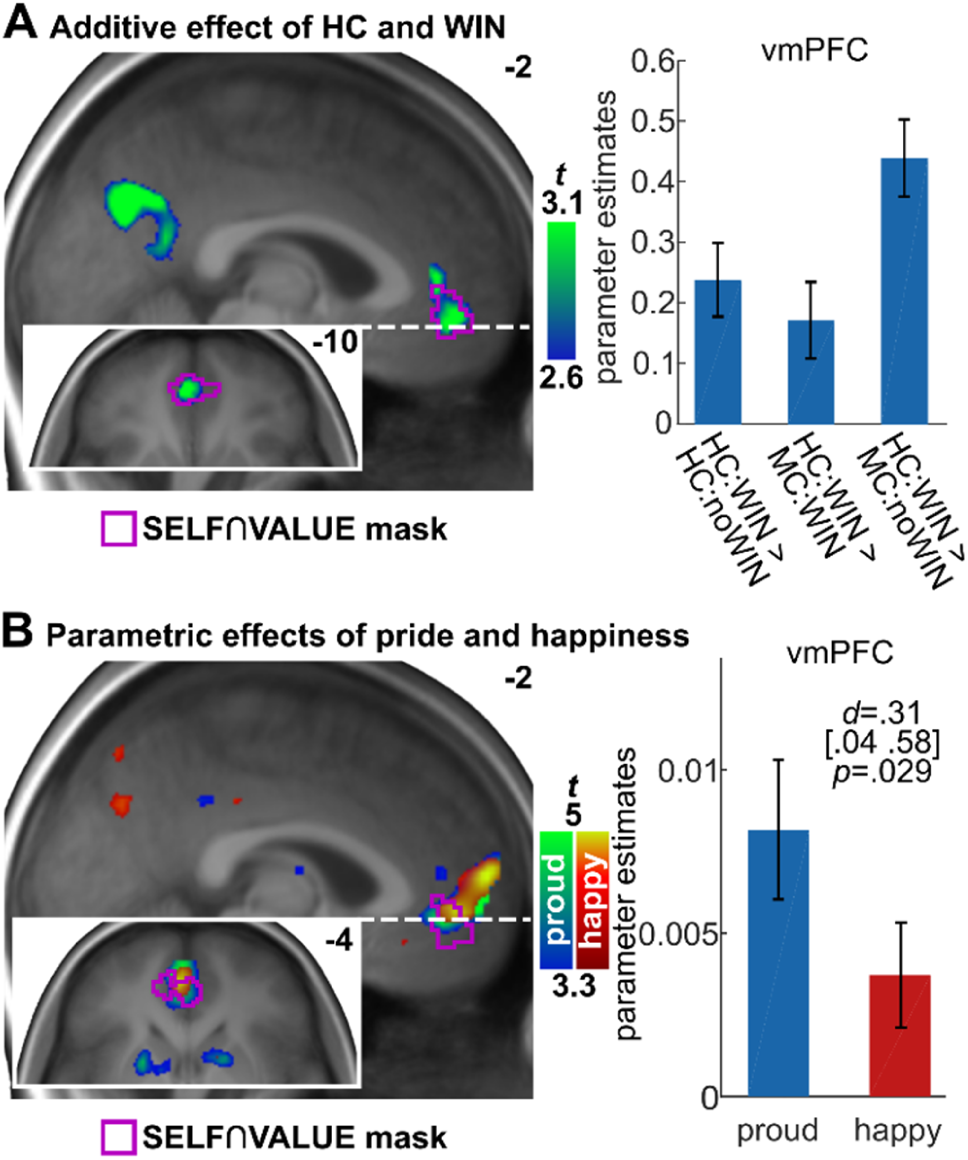
Additive effect of winning under high control and parametric effects of emotion ratings. **A** Activations show the specific effect of HC:WIN (i.e. 3-way conjunction [HC:WIN>HC:noWIN] ∩ [HC:WIN>MC:WIN] ∩ [HC:WIN>MC:noWIN], displayed at *p*<.005, uncorrected. The conjunction contrast survived small-volume FWE-correction at *p*<.05 inside the SELF∩VALUE mask in vmPFC. Bar plot represents differences of neural activation in vmPFC between HC:WIN and the other three outcomes. **B** Parametric modulation effect of happiness (red-yellow) and pride ratings (blue-green) at outcome presentation, displayed at *p*<.001, uncorrected for illustrative purposes. Bar plot shows group-level parameter estimates of the parametric modulations, averaged across all voxels inside the SELF∩VALUE mask. Violet outlines show SELF∩VALUE mask. Error bars are +/− 1 standard error of the mean. *d*=Cohen’s *d*. Values in brackets are lower and upper bound of 90% confidence interval for *d*.

#### Neural Correlates of self-evaluative Affect

We next aimed to link BOLD responses of task outcomes to intra-individual dynamics in the experiences of pride and happiness (figure 5). We found that within-subject dynamics in happiness were significantly correlated with neural activity within the VALUE­SELF mask at outcome presentation (16,2,−12, *t*(38)=4.55, *Z*=4.04, *k*=3, *p*=.022), as well as the SELF∩VALUE mask (left vmPFC; −2,48,−4, *t*(38)=4.60, *Z*=4.08, *k*=47, *p*=.002). Within-subject variability in pride ratings showed more extended significant effects in the SELF­VALUE mask (dorsal medial prefrontal cortex dmPFC; −2,56,2, *t*(38)=5.96, *Z*=4.98, *k*=122, *p*=.001) and the VALUE­SELF mask (dmPFC; −2,56,0, *t*(38)=6.15, *Z*=5.09, *k*=22, *p*<.001). In addition, variability in pride ratings correlated significantly with BOLD responses in the SELF∩VALUE mask (left vmPFC; −4,48,−6, *t*(38)=4.83, *Z*=4.24, *k*=94, *p*=.001; all *p*-values small-volume FWE-corrected; see Supplements 2). Notably, pride ratings covaried more strongly with neural activity in vmPFC than happiness ratings (paired-sample t-test on mean beta estimates for all voxels inside the SELF∩VALUE mask: *t*(38)=1.951, one-sided *p*=.029, *d*=.31, *CI*=[.04 .58]), indicating that neural activity in vmPFC is more strongly linked to affective experience when the causes of such experiences allow self-evaluation and might bear implications for one’s future behavior^27, 48^.

#### Dynamics of striatal Connectivity in the Context of Internal Control over Outcomes

Psychophysiological interaction (PPI) analysis showed that when task outcomes were received in the HC condition, functional connectivity increased between the left VS and left vmPFC (6,48,−14, *t*(37)=3.80; *Z*=3.47, *k*=48, *p*=.017; small-volume-corrected within the SELF∩VALUE mask). Though much more spatially extended in the CMS, this effect partly converged with the additive effects of success and control beliefs reported above. Beyond that, the left VS BOLD signal was more strongly associated with responses in the left angular gyrus (left AG) during outcome presentation in the HC as compared to the MC condition (−46, −70,38, *t*(37)=5.64; *Z*=4.76, *k*=24, *p*=.001 small-volume FWE-corrected inside the SELF­VALUE mask, surviving whole brain FWE-correction: −44,−70,40, *t*(37)=5.67, *Z*=4.78, *p*=.035). The equivalent analyses for the right VS showed significant effects in left superior frontal gyrus (−18,38,46, *t*(38)=4.59; *Z*=4.07, *k*=4, *p*=.028, small-volume FWE-corrected inside the SELF­VALUE mask), but not in vmPFC (4,38,−4, *t*(38)=3.23; *Z*=3.02, *p*=.069, small-volume FWE-corrected inside the SELF∩VALUE mask). At a more lenient exploratory threshold, for both left and right VS, we observed spatially extended connectivity changes in regions covered by the SELF­VALUE and SELF∩VALUE masks, such as the left AG, the dmPFC, ACC, and the vmPFC (*p*<.001; see Supplement 2). These findings align with the notion that outcome-related information processed in regions such as the ventral striatum^24^ are more strongly integrated with self-related representations in contexts with internal control, allowing to relate one’s own actions to ensuing outcomes.

Based on the assumption that stronger integration of self- and outcome-related processing should relate to more pronounced pride responses, we performed additional exploratory analyses on the connectivity dynamics induced by contexts of internal control beliefs. These showed that the observed increases of connectivity with the VS correlated significantly with the behaviorally measured pride response, at least in the left AG (*r*=.37, *p*=.032; dmPFC: *r*=.23, *p*=.080; precuneus: *r*=.23, *p* =.080, FDR-corrected one-sided *p*-values; figure 6, see methods for details). This implies that pride dynamics relate to changes in functional coupling of the VS with brain regions associated with self-referential processes. The equivalent analyses for the happiness response did not show any significant effects (left AG: *r*=.08, *p*=.582; dmPFC: *r*=.02, *p*=.582; precuneus: *r*=−.03, *p* =.582; FDR-corrected one-sided *p*-values). However, we note that these correlations did not differ significantly between pride and happiness (left AG: Hotelling’s *t*(35) = 1.84, *p*=.088; dmPFC: Hotelling’s *t*(35) = 1.29, *p*=.102; precuneus: Hotelling’s *t*(35) = 1.60, *p*=.088; FDR-corrected one-sided *p-*values). While general positive affect has recently been shown to relate to neural activity also elicited by surprising and uncontrollable task outcomes^33, 34^ these data demonstrate that positive affect is also considerably shaped by perceived control over task outcomes. Precisely, internal control beliefs increase self-relevance of task outcomes, driving activity in CMS and dynamics of self-related affect upon outcome reception^31, 53, 54^. These shifts in affective and neural dynamics effectively display as increased reports of pride experiences^55^, which relate to activity in the vmPFC as shown above and are assumed to bear implications for future behavior^31, 48^.

**Figure 6.**
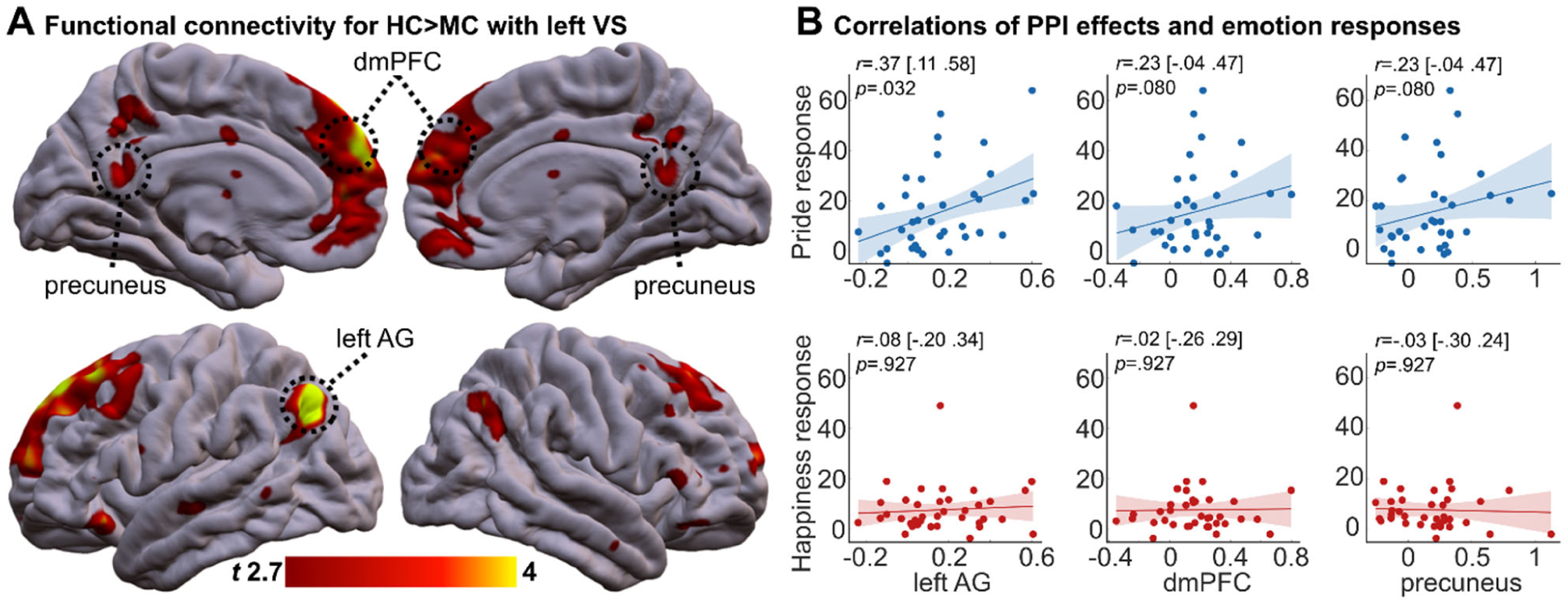
Increased functional connectivity with left VS in response to outcomes obtained under high subjectively perceived control tasks. **A** ROI analysis showed increased coupling between left VS and the left AG, the dmPFC and precuneus (small-volume FWE-corrected at *p*<.05). Effects in are displayed at *p*<.005, whole-brain, for illustrative purposes. See Supplement 2 for full connectivity coordinates. **B Top** Bivariate correlations between pride response (i.e. [HC:WIN - HC:noWIN] - [MC:WIN - MC:noWIN]) and estimates of functional connectivity from left AG, dmPFC and precuneus. Eigenvariates of clusters within the SELF¬VALUE mask consisting of voxels showing significantly increased functional connectivity with VS at *p* < .0005 (uncorrected) were computed for the three regions highlighted in A. *p-value*s are one sided and false-discovery-rate corrected. **Bottom** Analyses for happiness response equivalent to the ones performed for pride ratings. *r* = Pearson correlation coefficient. Values in brackets are 90% confidence intervals and shaded areas represent 95% confidence intervals.

### Study 3: Pride as an Affective Marker for the Subjective Value of Control

In study 3 we aimed to predict behavior based on affective experiences of pride in high control environments. We show that participants’ preference for controllable tasks is related to their pride response, even at a monetary cost. Subjects (*N*=50) completed a total of 64 trials of MC and HC (32 each), with pride ratings acquired after 40 outcome presentations. Following the task presented in study 2 (hereafter: the main task), we subsequently assessed whether subjects preferred HC or MC when different monetary amounts were offered. We employed an adaptive staircase algorithm^56^ thus varying the offers presented for HC and MC on every trial in a fashion allowing to identify each subject’s decision point. To increase the relevance of the task, subjects were informed that at random intervals some trials were going to be executed right after their choice. The chosen option, i.e. HC or MC with the associated potential payoffs, could then be played and the sum earned through successful task completion was awarded at the end of the experiment. Eight trials were pseudorandomly selected and presented to the subjects after the respective choice, however without the presentation of task outcomes.

Confirming studies 1 and 2, subjects experienced increased internal control in HC than in MC (*t*(49)=11.35, *p*<.001, *d*=1.61, 90% *CI*=[1.25; 1.95]). Again, pride reactivity to outcomes differed between HC and MC (condition*outcome valence: *F*(1,49)=35.98, *p*<.001, 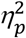=.42; figure 7). Specifically, pride was rated higher for WIN than for noWIN outcomes (MC: *t*(49)=7.15, one-sided *p*<.001; HC: *t*(49)=12.37, one-sided *p*<.001, Bonferroni-corrected for multiple comparisons). In addition, the difference in affect ratings between WIN and noWIN outcomes was larger in the HC condition than in the MC condition (*t*(49)=6.00, one-sided *p*<.001; see Supplement 1 – table S3).

**Figure 7.**
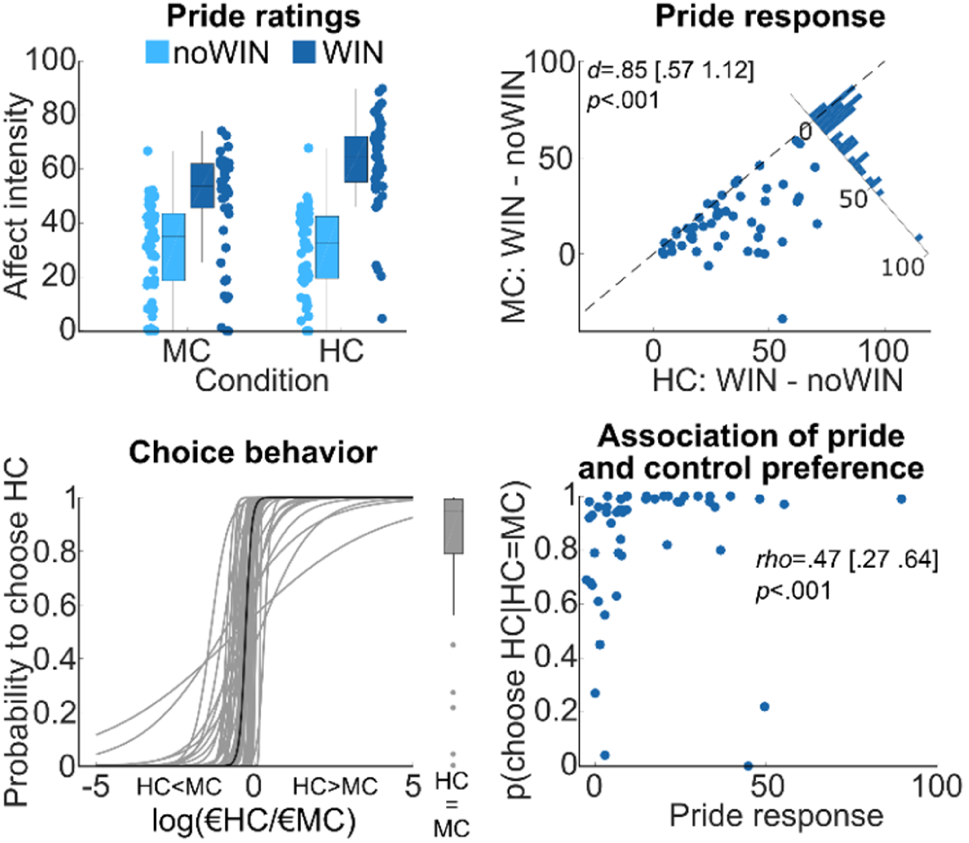
The experience of pride in response to high control affects subjective value of choices. **Top left** Pride ratings for study 3, separate for high and low control, as well as outcome valence. **Top right** Pride reactivity to WIN vs noWIN outcomes, separately for HC (x-axis) and MC (y-axis). Histogram depicts difference between HC and MC. *d*=Cohen’s *d*. Values in brackets represent lower and upper bound of 90% confidence interval for *d*. **Bottom left** Probability to choose HC in choice task of study 3 as a logistic function of log-transformed relative option values (log(€HC/€MC)). More positive values on horizontal axis indicate higher monetary value of HC relative to MC. Thin grey lines represent individual participants’ choice functions. Thick black line represents hypothetical choice behavior based on sample medians for *β*_0_ and *β*_1_. Boxplot represents participant’ probabilities to choose HC, given that both options have equal values. **Bottom right** Relationship of pride response and choice behavior in study 3. *rho*=Spearman’s rho. Values in brackets are lower and upper bound of 90% confidence interval for *rho*.

In the choice task, the mean percentage of choices in favor of the HC option was 58.1% (*SD*=10.2%), which was significantly higher than chance (*t*(49)=5.62, one-sided *p*<.001). The mean value of the chosen MC options was 1.19€ (*SD*=.09€) and that of the chosen HC options was 1.03€ (*SD*=.07€; paired *t*-test: *t*(49)=8.20, one-sided *p*<.001), indicating that participants had a preference for the HC task, despite having a 13.4% smaller payoff than for the MC options.

In order to better characterize the choice behavior, we assessed the model parameters of a logistic choice model fit to each participant’s data (see methods). Across participants, the mean value of the models intercept *β*_0_ was significantly negative (*M*=−3.49, *SD*=4.18, *t*(49)=−6.00, one-sided *p*<.001) indicating that participants preferred the HC option over the MC option, if both options offered equal monetary payoffs. Similarly, *β*_1_ was significantly negative (*M*=−17.21, *SD*=19.67; *t*(49)=−6.19, one-sided *p*<.001), showing that across participants, as expected, the probability for choosing HC increased with higher monetary gain for HC relative to MC (figure 7). Since *β*_0_ is directly convertible into the probability to select HC given equal monetary payoffs of the two options, we will base the following analyses on this more intuitive measure. More specifically, if the nominal monetary offer for both options was equal, there was a strong preference for HC, with over 90% of the participants preferring the controllable over the less controllable option (*MD*=95.1%, *IQR*=20.48, figure 7), which was significantly above 50% (Wilcoxon signed-rank *W*=1182, one-sided *p*<.001). Importantly, the subjects’ probability to choose HC given equal expected values for HC and MC remained significantly above 50% (*MD*=65.98%, *IQR*=50.48, Wilcoxon signed-rank *W*=863, one-sided *p*=.015) even when considering differences in expected outcome probabilities, by controlling for differences in subjectively perceived winning histories, that deviated from the objectively presented outcome rates of 50% for the two tasks (see Methods).

A further goal of study 3 was to demonstrate that participants’ preference for HC over MC varies in dependence of the affective dynamics during the main task. As expected, individuals who had a more positive pride response showed a stronger preference for HC in the choice task, given equal values of the two choice options (Spearman’s *rho=*.47, one-sided *p*<.001; figure 7). This association remained significant even after controlling for various control variables such as expended effort or perceived winning probabilities (see Supplement 1 – table S6). This extends previous results showing that mere choice is preferable to not having choices by highlighting the notion of instrumental control over the outcomes for building preferences^26^, and emphasizing that self-related positive affect is a relevant factor for motivated behavior^28^.

### Studies 1 – 3: Interindividual Variability in Control Beliefs Predicts Pride Response

Aggregating data from all 129 participants, we finally tested whether interindividual differences in the degree to which the task could stimulate control beliefs were associated with self-related affective experiences. Across the three studies, participants who perceived more control in HC compared to MC also showed stronger pride responses (Spearman’s *rho*=.25, one-sided *p*=.003; figure 8), supporting the notion of how individual differences in the experience of internal control drive self-conscious positive affect^27, 31^.

**Figure 8.**
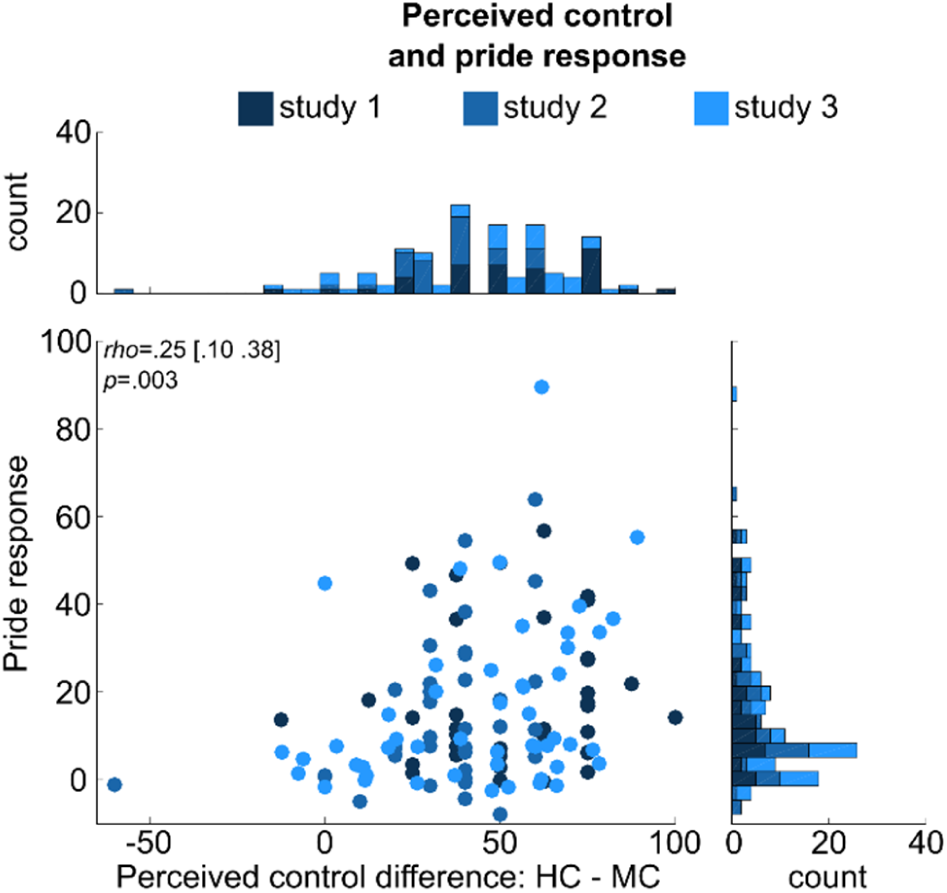
Association of pride response and individual differences in the experience of control. The scatterplot depicts the relationship between internal control beliefs and pride response across all three studies. Dark blue indicates data point belongs to study 1, intermediate blue to study 2, and light blue to study 3. Raw controllability ratings were transformed to a range between 0 and 100, across internal control conditions, before computing the difference between HC and MC in each study. For study 1, raw control and pride ratings were averaged across LC and MC. The pride response is the interaction term for pride ([HC:WIN–HC:noWIN] - [MC:WIN–MC:noWIN]). Histograms depict counts for difference in perceived control (top), and pride response (right). Values in brackets represent lower and upper bounds of the 90% confidence interval for

## Discussion

By employing a novel experimental paradigm, we demonstrate that task environments with internal control beliefs entail unique affective consequences with associated shifts in neural processing. Precisely, tasks offering internal control allow self-attribution of outcomes. Thereby, such contexts differentiate self-conscious positive affect – i.e. pride – from more unspecific positive affect – i.e. happiness – when receiving task outcomes. Using functional magnetic resonance imaging, we furthermore characterize brain regions involved in altered valuation of outcomes achieved in controllable tasks. In the task environment in which outcomes more strongly depend on own-contributions we see increased activity in brain regions of the cortical midline, such as the medial prefrontal cortex and precuneus, that are associated with self-related processing^53, 54^. Here, the vmPFC seems to play a key role in tracking both success as well as internal control. In addition, activity in these regions is more strongly related to functional responses of brain regions processing reward information, i.e. ventral striatum, when outcomes depend on one’s own behavioral performance. In a final step, we highlight the subjective value of the emerging self-related affect, as people discount monetary gains for tasks offering internal control in dependence of their pride response during task execution.

Overall, our findings provide evidence for the link between internal control beliefs and the affective reaction of pride. This concords with psychological theories of self-conscious affect, stating that internal control allows attributing task outcomes to internal causes, such as one’s abilities^1, 27^. Self-evaluative processes are crucial to the understanding of self-conscious affect like pride^27^, differentiating them from other affect constructs such as happiness. While positive affect was generally increased in response to successes under high control this was particularly the case for pride responses and less so for participants’ reports of happiness which conceptually does not necessarily includes processes of evaluating one’s self-worth^27, 57^. Since in potentially controllable contexts outcomes are informative about one’s abilities, participants can use these outcomes to infer to what extent they succeed at actually exerting control. Regarding affective responses, this was reflected by the fact that pride was most prominent in the task allowing the highest levels of internal control.

Importantly, individuals with stronger control-dependent pride responses preferred to partake in the task offering internal control over another, less controllable one. Concordant with earlier findings^20, 28, 58^, and substantiating the claim that self-conscious affect is central in the guidance of future behavior^27^, this finding underlines the motivational effects of pride^30^. Strikingly, preferring to choose an opportunity to exert control outweighed a monetary detriment entailed by this choice behavior. Collectively, this indicates that the development of behavioral preferences is partially driven by the extent to which an individual perceives contexts to be controllable and how succeeding in such contexts manifests in the experience of self-related affect, such as pride. That is, the experience of mastering a task might elicit feelings of pride and competence that increase the probability of choosing to perform the same or similar behaviors later on, dovetailing nicely with theories linking experiences of competence to intrinsic motivation^1^.

Guided by automated meta-analysis^52^, we show that these effects of self-attribution on outcome processing are mediated via two distinctive, but connected brain networks associated with the processing of value and self-related information. Outcomes from tasks with high control were associated with activity in brain regions linked to self-related information processing, such as CMS^45, 46^, since outcomes in controllable tasks are informative about one’s abilities and thus inherently self-relevant. This is an important amendment to previous evidence and underscores that control beliefs need to be understood beyond having a choice or not in order to understand how subjects develop self-esteem^32^ and concepts of their own capabilities in acting on the environment^1^. Our findings thus are in line with other studies showing that ACC is implicated in monitoring self-initiated actions^59^ or vmPFC being involved in outcome monitoring, especially when events are relevant to the individual^45^. However, while paralleling these earlier findings, our data also provide novel insights by relating these processes to the concept of self-conscious affect. First, in the condition eliciting the strongest pride, i.e. when receiving outcomes indicating success in a controllable task, the vmPFC was significantly more active than in any other condition. Second, in contexts of internal control beliefs, regions associated with processing self-related information such as left AG, vmPFC, and dmPFC, were more strongly coupled to VS, which is implicated in the processing of reward information^24^, and these effects correlated with behaviorally measured pride responses across subjects. Third, intraindividual variability in pride covaried more strongly than happiness with neural activity in vmPFC. Collectively, these findings support the notion that self-related positive affect might depend on the integration of outcome information with self-representations, mediated via regions of the cortical midline and AG, mirroring findings of studies investigating other types of self-conscious affect^60^.

Regarding the relationship of affect and neural processing, Rutledge and colleagues^33^ more recently linked moment-to-moment variability of happiness to errors in reward prediction, which in turn is associated with activity in the VS. In a follow-up study, they highlighted the importance of investigating “instrumental control” for a deeper understanding of well-being^34^. Our study takes a first step in this direction by manipulating internal control beliefs, underlining its importance for differentiated affective experience and downstream motivational effects that manifest in behavioral preferences. However, our study design does not allow explicit modeling of reward expectations and associated reward prediction errors. While recent work suggests how control beliefs and prediction error processing might interact^19, 26, 37^, explicit neuroscientific investigations of their interplay are lacking. Thus, future studies should directly test how prediction error signaling changes in controllable environments, how this relates to affective dynamics, and how these processes are mediated neurally^34, 61^.

Our findings replicate and extend previous work on the value of control^18, 20^ by demonstrating that participants preferred a controllable task over an uncontrollable one, reflecting differences in subjective value that are likely processed by vmPFC^15^. This is supported by previous studies showing activation differences in vmPFC for the anticipation of choice vs. non-choice^20^. However, the core feature of our study was a condition supplying participants with an internal model of how their behavior maps to outcomes, 22 extending the concept of control beliefs beyond merely having a choice^26^. Rather, the high control condition in our task suggested that the subjects’ performance determined whether or not a goal was reached. This condition yielded the strongest affective reactivity to outcomes, which was reflected in activity changes in the vmPFC within subjects and increased functional coupling of the vmPFC with VS. Hence, vmPFC might serve the integration of outcome information with representations of one’s abilities^50, 54, 62^, potentially driving differences in behavioral preferences for subjectively more or less controllable tasks^15, 26, 42^. Future research could follow this path by assessing directly how control-dependent outcome valuation translates into behavioral preferences in subsequent choice situations and how this might be mediated neurally by processing of subjective value in the vmPFC.

Pride experiences in response to outcomes that can be attributed to the self foster self-esteem. Thereby they effectively shape meta-cognitive beliefs about oneself and estimates of one’s social status^32, 63^. A recent study investigated updating of state self-esteem in response to evaluation through others^51^ and found these updates to be positively correlated with activity in vmPFC. Comparably, in a study in which participants were asked to imagine or remember pride-associated events, significant activations in the medial prefrontal cortex were found that extended into vmPFC^64^. Furthermore, two recent studies have supported this role of vmPFC in the updating of self-related beliefs^50, 65^. In this perspective, the vmPFC activation and affective dynamics we observed in response to outcomes from controllable tasks might reflect correlates of short-term updating of beliefs about one’s abilities. If subjects experience pride this might contribute to updating their concepts of their own capabilities in a specific environment. In the long run, such positive self-evaluations support self-esteem and help individuals with developing an understanding of their worthiness^66^.

From a broader perspective the findings described here have implications for learned helplessness and psychiatric conditions, such as major depressive disorder^13, 42^. Our results emphasize that lower internal control beliefs are associated with reduced attribution of positive outcomes to the self, relating to decreased pride experiences. Further, these reductions in self-related affective responses align with 23 diminished preferences for tasks offering opportunities for exerting control. These dependencies might contribute to reduced self-esteem in the long run^32^ and reduced motivation to show goal-directed actions^1, 13^. Such processes could therefore causally relate to depressive episodes, which are centrally defined by negative views of the self and lack of motivation^14^. In contrast to states of reduced internal control, other psychiatric phenomena such as pathological gambling are characterized by an increase in internal control beliefs, even in contexts offering no evidence for actions being linked to outcomes by ways other than chance^67–69^. In this regard, our data hint at a possible role of affective reactions to (un)controllable events in mediating the development of gambling addictions^70^. Individuals who perceive control over objectively uncontrollable events might experience positive self-related affect once a desired outcome – such as “cracking” a one-armed bandit – is obtained, and might thus be at greater risk of developing gambling addictions. Therefore, the findings presented here provide a first hint for future clinical research to refocus on perturbed beliefs of control over the environment to understand affective and motivational symptoms as well as altered neural dynamics associated with psychiatric conditions.

In conclusion, the three studies provide unequivocal evidence that internal control beliefs impact the valuation of outcomes, resulting in entangled positive self-conscious affect and behavioral preference for environments with greater perceived control. While based on a classical understanding of control beliefs^4, 6, 7^, the present studies add to our understanding of how outcomes that have been achieved through one’s own actions recruit neural systems involved in computations of reward value and self-relatedness^54^. Increased interactions of the ventral striatum and cortical midline structures, as well as responses of the vmPFC that track both shifts in outcome processing under internal control beliefs and self-conscious positive affect thereby provide first links to how self-attribution of desirable events drives motivation and future behavior^26^. On a more general note, our studies underline the need for the neurosciences to consider how behavior and subjective experiences are shaped by beliefs individuals hold, that regard how their actions relate to events in the world they live in and interact with. Taking further steps in this direction promises to deepen our understanding of affect, motivation, and underlying neural dynamics with important implications for scientific models of behavioral control, general well-being, and psychiatry^1, 2, 26, 37^.

## Material and Methods

### Participants

A total of 154 participants took part in the three studies (mean age: 23.41 years; SD: 3.41; range, 18-37; 77 females). All experimental designs were approved by the ethics committee of the University of Lübeck, Germany (AZ 16-133). Participants had to be right-handed, speak fluent German, have no deficits in color vision, normal or corrected-to-normal vision, and no pre-existing psychiatric or neurological conditions. Psychology students (study 1: in the second or later year of university) were not invited. In study 2, contraindications for fMRI scanning (e.g. pregnancy, metal parts inside the body) were additional exclusion criteria. All participants gave written informed consent prior to participation in the study, were debriefed after completion of the experimental session, and received monetary compensation or partial course credit for participation. See table 1 for details on the individual samples.

**Table 1.**
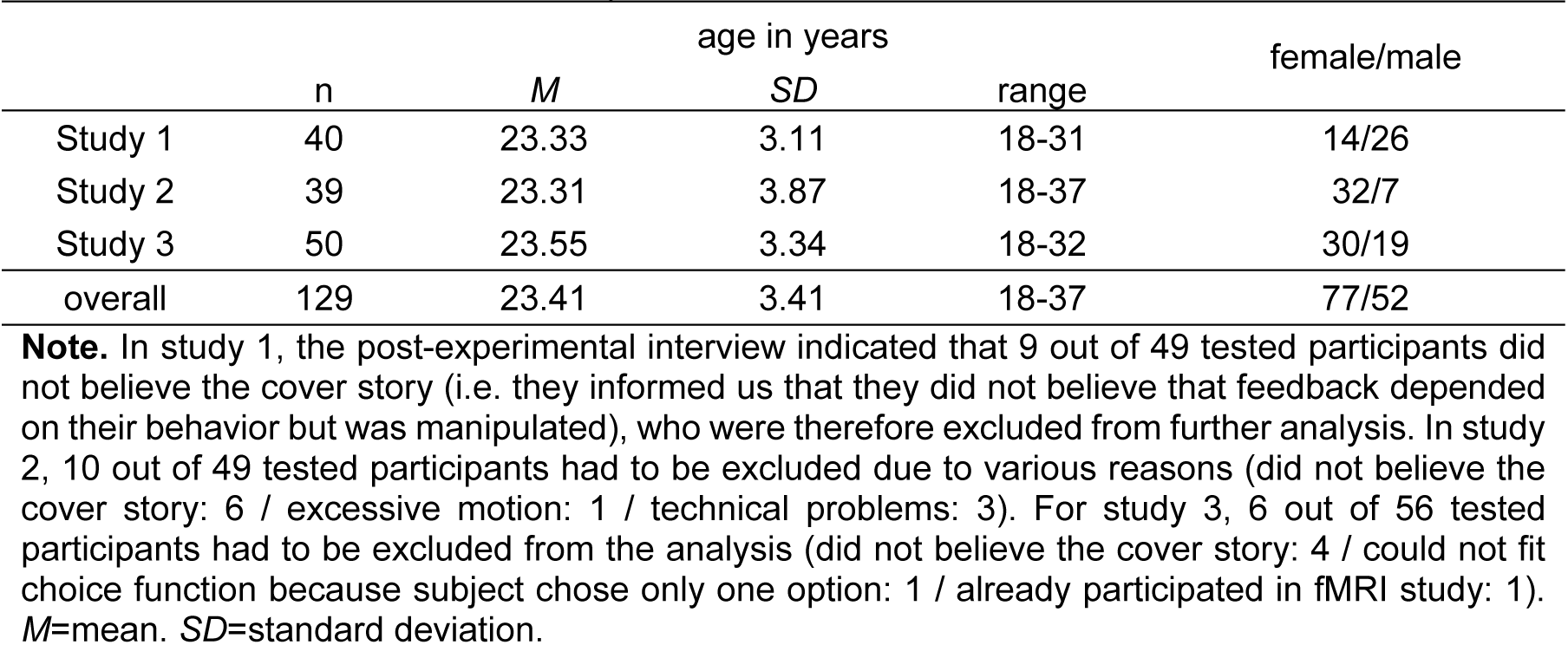
Descriptive statistics for study samples

|  | n | *M* | *SD* | age in years<br>range | female/male |
| --- | --- | --- | --- | --- | --- |
| Study 1 | 40 | 23.33 | 3.11 | 18-31 | 14/26 |
| Study 2 | 39 | 23.31 | 3.87 | 18-37 | 32/7 |
| Study 3 | 50 | 23.55 | 3.34 | 18-32 | 30/19 |
| overall | 129 | 23.41 | 3.41 | 18-37 | 77/52 |

**Note.** In study 1, the post-experimental interview indicated that 9 out of 49 tested participants did not believe the cover story (i.e. they informed us that they did not believe that feedback depended on their behavior but was manipulated), who were therefore excluded from further analysis. In study 2, 10 out of 49 tested participants had to be excluded due to various reasons (did not believe the cover story: 6 / excessive motion: 1 / technical problems: 3). For study 3, 6 out of 56 tested participants had to be excluded from the analysis (did not believe the cover story: 4 / could not fit choice function because subject chose only one option: 1 / already participated in fMRI study: 1). *M*=mean. *SD*=standard deviation.

### Design and Procedure

#### Description of the main experimental task

In the main experimental task we manipulated internal control beliefs over task outcomes across three levels (low, medium and high control; in the following LC, MC and HC). The tile task displayed a grid-pattern of two halves of turquoise and yellow squares surrounding a single grey square in the center (figure 1). On each trial, orientation of halves (left-right or upper-lower arrangement) and brightness of squares within each half was pseudo-randomly defined. On each trial, three, four or five squares on each half (targets) were assigned a brightness that was considerably higher than the remaining ones (distractors). Targets never appeared side by side. Crucially, while there were random differences in brightness of all distractor squares, the brightness of target squares was kept near identical rendering a definite identification of the brightest square within one half impossible. The brightness of the grey square in the center remained constant across the experiment.

Perceived controllability of task outcomes was manipulated in a within-participants design. In the low control (LC) condition, participants were instructed to *click* on the central grey square. In 50% of trials clicking the square was pseudo-randomly followed by a win and in the other 50% by a no win outcome (in the following: WIN and noWIN). In the medium control (MC) condition, participants were instructed to *choose* one color (i.e. yellow or turquoise) by clicking on any square of that color. Again, in 50% of trials choosing a half was pseudo-randomly followed by either a WIN or a noWIN outcome. In the high control (HC) condition, participants were instructed to *identify* the brightest square of either the yellow or turquoise half of the grid. Again, in 50% of trials the identification of the putatively brightest square was pseudo-randomly followed by either a WIN or a noWIN outcome.

In each study, given that no mistakes were made (i.e., response was fast enough, and participant did not click on background instead of a square, and a square of the correct color was clicked in the HC condition), an equal amount of trials was presented for every combination of outcome valence (WIN, noWIN) and control (LC, MC, HC in study 1; MC, HC in studies 2 & 3).The number of ratings was equally distributed across combinations of outcome valence and internal control condition and the same for pride and happiness. The level of internal control regarding each condition (HC, MC, and LC; “How strongly could you influence / control the outcome in the different tasks?”, ranging from “very strongly” to “not at all”; see below) was rated after the main task using 5-point Likert-type scales in studies 1 & 2, while a continuous horizontal rating scale was used in study 3.

All subjects in a given study were tested in the same laboratory room and lighting conditions were held constant by lowering the window blinds and setting the dimmable electric lights to the same level. Brightness of the 3-5 brightest squares was randomly set on each trial within these pre-defined limits in studies 1 & 2. In study 3, instead of creating a unique display by setting the previously described parameters randomly on each trial, a predefined pool of 100 grid stimuli was used, which was shuffled and therefore randomly assigned to each condition for each subject.

In each study, presentation of WIN and noWIN outcomes was manipulated, so that the overall probability of receiving each outcome would be fixed to 50% for every condition. In case a square was clicked that was not defined as belonging to the set of brightest squares on a given HC trial, a noWIN was presented in order to maintain credibility of the task. The predefined sequence of WIN and noWIN outcomes was designed in a way that no more than four WIN or noWIN trials appeared consecutively across conditions, and that no more than two WIN or noWIN trials appeared consecutively within conditions. Additionally, no condition appeared more than two times in a row. This was done in order to maintain an expectation of receiving a WIN (noWIN) outcome close to 50% across the course of the task. All experimental stimuli were presented using the Psychophysics Toolbox^71^ for Matlab (Mathworks Inc, Natick, MA). Post-experimental interviews were administered using computerized questionnaires presented in Sosci Survey (Sosci Survey GmbH, München, Germany).

#### Study 1: Dynamics of self-evaluative positive Affect

Prior to the main experiment, and after a short demonstration of the three task conditions, participants completed a practice session (see below) so that they could learn a) the timing of the task, and b) that they would receive approximately 50% WIN and 50% noWIN feedbacks in all three conditions.

After the practice session and prior to the main task, participants completed two trials of every condition including one WIN and one noWIN outcome each, and ratings of pride and happiness in order to learn the timing of the complete trial structure. Subsequently, participants performed a total of 90 trials (figure 1B) of the experimental task. The sequence of conditions and outcome valence in each condition 28 was predefined, such that no condition appeared more than twice in a row. Additionally, no more than four consecutive WIN or noWIN trials were presented across conditions, and no more than two repetitions of the same outcome valence appeared within conditions. Ratings of pride and happiness were displayed side-by-side after each trial and visualized as thermometers initiated at 0 and ranging to 100. Participants could rate their subjective feelings of pride and happiness in any order by clicking on the corresponding levels of the two thermometers and confirming their ratings using a *done*-button on the bottom of the screen.

#### Study 2: Neural Foundations of Outcome Valuation in the Context of internal Control

The procedure in the study 2 largely followed that of the study 1, with a few changes. First, the LC condition was not presented in the fMRI in order to reduce scanning time and to perform a more conservative comparison between the MC and HC conditions that both had a component of choice. A total of 80 trials were presented, followed by a rating of either pride or happiness (figure 1B) on each or every second trial. If no mistakes were committed, this yielded a total of 24 ratings each for pride and happiness, 12 for each condition, and 6 for every combination of WIN/noWIN and condition. For approximately half of the participants, the trials on which pride and happiness were rated were switched. In case a mistake was committed, both the feedback and the rating on that trial were replaced by a warning and a prolonged inter-trial-interval (ITI) until the planned onset time for the next trial was reached.

After a short explanation of the task outside the scanner, the practice session in the fMRI study was done inside the fMRI scanner and followed the same structure as in the study 1, as did the post-experimental interview and debriefing. Participants completed all tasks with an fMRI-compatible computer mouse (NAtA TECHNOLOGIES, Coquitlam, Canada) and their right arm rested on a tilted mousepad supported by an adjustable armrest. ITIs were pseudo-randomly jittered between 3-5 seconds during which a fixation cross was presented in the center of the screen. Following a 1 second cue, the task was presented for 4.75 seconds and after a 4 second delay, the outcome was presented for 3 seconds. On each or every second trial, participants were asked to rate within 4 seconds either how proud or how happy they felt regarding the preceding outcome. Ratings were given on a horizontal rating scale initiated at an intermediate position (“somewhat”) and ranged from “not at all” to “very” (figure 1), and converted to values ranging from 0 through 100.

#### Study 3: Pride as an Affective Marker for the Subjective Value of Control

Study 3 consisted of two behavioral paradigms (figure 1B). After an initial demonstration of the task and a practice session, 64 trials of the main task were presented, and pride ratings were collected on 40 pseudo-randomly selected trials. Thereafter, participants completed a choice task in which on each of 96 trials they were given a choice between performing either the MC or the HC condition. Importantly, each choice option was associated with varying amounts of money (see below for details). For instance, a subject could have the choice between participating in the MC condition for a potential gain of 1 € or, alternatively, participate in the HC condition for a potential gain of 0.9 €. On eight pseudo-randomly selected trials, participants performed the chosen task, and were informed that after the experiment they would receive the sum of all successfully completed trials. No outcomes were presented during this part of the experiment to avoid influencing the participants’ expectation of succeeding.

The algorithm we used to determine the potential monetary gains of the choice options on each trial followed the logic of an adaptive staircase algorithm for the detection of perceptual detection thresholds as used in psychophysics^56^. Specifically, on each trial, one of the two choice options was selected as the *reference* and assigned a value of 1€, while the other option was assigned another value that was either higher or lower than the reference value. E.g., if this *comparison value* was higher than the reference value on a given trial and the participant picked the higher value, the comparison value was decreased by a predefined fraction in the following trials until the participant changed her decision and chose the option with the reference value. In this fashion we aimed to find the point at which each individual participant valued both choice options equally. A description of the staircase-algorithm we employed is detailed below.

Between the main task and the choice task, participants gave a number of ratings regarding their experience of the task. These ratings included the participants’ estimate of the percentage of WIN and noWIN outcomes they had received during the different conditions in the main task, or they thought they would receive if they continued to perform the same conditions in the future.

#### Practice Task (studies 1, 2, & 3)

The practice task preceding the main experiment was conducted in order to minimize differences in outcome expectations between the three conditions. During the practice session, participants completed 15 trials of each condition without any feedback during the first twelve trials. Instead, subjects received manipulated feedback during the last three trials of the different conditions, indicating that they had “won” (LC), “chosen correctly” (MC), or “found the brightest square” (HC) in approximately 50% of the preceding trials. Precisely, there were three sets of feedback percentages (46, 50, 54 / 47, 50, 53 / 48, 50, 52), that were randomly assigned to the task condition based on the subject ID and the order of each set was randomized for each condition and every subject. In case a subject made too many mistakes (i.e. responded too slowly, clicked on the background, clicked a square of the incorrect color in the HC condition, or did not click on the grey square in the LC condition on more than ¼ of the completed practice trials) after the first eleven practice trials of a condition, the subject was informed about this by an on-screen message and the respective practice block was repeated. Before the start of the practice session, subjects were informed that depending on their performance in the practice blocks, they would be assigned to one of three groups gaining either 10, 15 or 20 €-cents on each correct trial of the main task and that the better they performed the more money they could gain during the main task. After the practice session, all subjects were informed that they had been assigned to the 20-cent group (irrespective of their performance).

#### Description of Ratings

In study 1, ratings of both pride and happiness were administered after every outcome, using thermometer-like rating scales, ranging from 0 through 100, and initialized at 0 (see figure 1). Subjects could decide which affect to rate first and completed their rating by clicking a *done*-button. In study 2, either pride or happiness ratings were administered after a given trial, with a total of 24 ratings for each affect (6 for each combination of outcome valence (WIN, noWIN) and control (MC, HC). For 20 out of the 39 subjects included in the analyses in study 2, the predefined assignment of pride and happiness ratings to a given outcome was switched. In study 3, only pride was rated after a total of 40 trials, equally distributed over all combinations of outcome valence and control. In studies 2 and 3 ratings were administered using horizontal rating bars, asking subjects how strongly they felt a given affect with regard to the preceding outcome. Scales ranged from *not at all* through *somewhat* to *very* and were initialized at *somewhat*, i.e. a neutral position.

Ratings of internal control beliefs in studies 1 and 2 were assessed using a paper-pencil 5-point Likert scale asking participants “How strongly could you influence the outcome in the different tasks?”, with the different options being: “very strongly”, “strongly”, “neutral”, “weakly”, “very weakly”. In study 3, two items were used to assess control beliefs, with the first item asking the same question as in studies 1 and 2. The second item asked “How strongly could you control the outcome in the different tasks?”. In study 3, a horizontal rating bar ranging from “very weakly” through “somewhat” to “very strongly” was displayed after the main task, similar to the affect ratings and initialized at a neutral position (i.e. “somewhat”). In study 3, ratings from both items were averaged for each condition in order to measure internal control beliefs.

In addition to ratings measuring control beliefs, after the main task in study 3, a set of additional questions was asked – separately for MC and HC – that was later used to statistically control the relationship between the pride response and choice behavior (Supplement 1 – tables S5 & S6). All questions were asked using a horizontal rating bar like the ones used for the affect ratings and the control ratings in study 3 and all answers were initialized at a neutral position and recoded to range from 0 to 100.

#### Choice Task (Study 3)

The algorithm we used to determine the potential monetary gains of the choice options on each trial followed the logic of an adaptive staircase algorithm for the detection of perceptual detection thresholds as used in psychophysics^56^. More specifically, on each trial, one of the two choice options was selected as the *reference* and assigned a value of 1€, while the other option was 32 assigned another value that was either higher or lower than the reference value, called the comparison value. This comparison value was changed on each trial by a certain step size in a specific manner detailed below.

In the beginning of the task, two step sizes were used: 33 cents and 17 cents. Additionally, the comparison value could either be smaller or larger than the reference value and was initially calculated by 3*step size, giving four sets of initial comparison values: 1.99€, 1.51€, 0.49€, 0.01€. Since either HC or MC could be the reference value on a given trial, we obtained a total of 8 initial starting values, defining 8 so-called comparison sets.

The task was programmed to contain four phases of 24 trials each that were not explicitly conveyed to the participants. In the first phase, only those comparison sets were used that started with a step size of 17 cents (i.e. comparison values starting at 1.51€ or 0.49€). In the second phase, two thirds of the trials belonged to these sets, while one third were trials starting with comparison values of 1.99€ or 0.01€ (i.e. step sizes of 33 cents). In the third phase of the task, two thirds of the trials belonged to the comparison sets with a step size of 33 cents and all of the trials in the fourth phase belonged to this set.

In each phase, 2 trials were randomly selected to be actually played by the subject (being HC or MC, depending on the subject’s choice), without showing an outcome. Subjects were informed that in the end, they would receive the amount gained from those trials on which they succeeded (i.e. found the brightest square in HC; selected the correct color in MC). However, they truly received 50% of the amounts they chose and played for during the decision task.

If on a given trial the comparison value (e.g. HC = 1.99€) was higher than the reference value (e.g. MC = 1€) and the subject picked the choice option with the (higher) comparison value, the comparison value was decreased by the current step size (e.g. 33 cents) in the following trials of the same comparison set until the subject changed her decision and picked the option with the reference value. As an example, the MC condition could be defined as the reference (1 €), while the initial comparison value of HC could be 1.99 €. Choosing HC for 1.99 € would lead to a decrease of the comparison value to 1.66 € on the next 33 trial of this comparison set. The comparison value would be decreasing as long as HC is preferred over MC. At some point, e.g. when HC has a value of .67 €, the subject might prefer MC (having a value of 1€). Now, the comparison value is *increased* until the subject’s decision changes a second time, and the comparison value starts to decrease again, this time by a smaller fraction than before (*previous step size*/1.25), therefore iteratively approximating the point at which the subjective value of both options are subjectively equal for this participant. On each trial, a random amount between −5 and 5 cents was added to the values in order to decrease monotony of the task. The highest offered value was 1.99€ and the lowest was 0.01€.

#### FMRI Data Acquisition

Participants were scanned using a 3T Siemens MAGENTOM Skyra scanner (Siemens, München, Germany) at the Center of Brain, Behavior, and Metabolism (CBBM) at the University of Lübeck, Germany. After four dummy-scans allowing for equilibration of T1 saturation effects, 768 volumes with 39 near-axial slices in ascending order were acquired for each participant using echo-planar-imaging (voxel size=3*3*3 mm, 68*68 matrix, 20% interslice-gap, TR=2000 ms, TE=25 ms, 90° FA, iPAT=2). In addition, a high-resolution anatomical T1 image was acquired that was used for coregistration (voxel size=1*1*1 mm, 256*256 matrix, TR=1900 ms, TE=2.44 ms, 9°FA).

### Statistical Analyses

The statistical analyses of subject ratings were performed using JASP version 0.9^73^. MRI data were analyzed using SPM12^74^, and the logistic choice models in study 3 were fitted using Matlab R2016a. The comparison of correlations between PPI-effects and affective responses in study 2, as well as FDR-correction of *p*-values regarding these correlations were performed in R^75^.

#### Manipulation Checks

In order to verify that subjects could not reliably detect the brightest square, we computed how many times in the entire experiment the brightest square of a given color would be found given a subject performed at chance level. For instance, if in a given trial three squares were defined to be the brightest squares, there was a chance of 1/3 to find the brightest square. We computed the level of chance performance for each subject individually, accounting for the fact that by chance, the 34 number of trials in which 3, 4, or 5 squares were the brightest could vary between subjects (due to the design of the experimental task). We performed paired t-tests in each study, comparing the number of times each subject found the brightest square with the level of chance performance. These analyses indicated, that in no study performance exceeded chance level. In fact, in every study, subjects performed significantly worse than chance (study 1: *t*(39) = −33.931, *p* < .001; study 2: *t*(38) = −2.311, *p* = .026; study 3: *t*(49) = −2.762, *p* = .008).

The objective rates of receiving WIN and noWIN outcomes did not deviate significantly from 50% in any study (study 1: LC: *M*=50.45%, *SD*=1.78, *t*(39)=1.601, two-sided *p*=.117, *MC*: *M*=50.26%, *SD*=1.29, *t*(39)=1.272, two-sided *p*=.211, HC: *M*=50.01%, *SD*=2.20, *t*(39)=0.030, two-sided *p*=.977; study 2: MC: *M*=50.10%, *SD*=1.14, *t*(38)=0.556, two-sided *p*=.581, HC: *M*=49.88%, *SD*=1.70, *t*(38)=−0.437, two-sided *p*=.664; study 3: MC: *M*=50.55%, *SD*=1.42, *t*(49)=0.275, two-sided *p*=.784, HC: *M*=50.00%, *SD*=1.15, *t*(49)=0.0005, two-sided *p*=.999), nor were there any significant differences in outcome rates between conditions (study 1: *F*(2,78)=0.572, *p*=.567; study 2: *t*(38)=−0.703, two-sided *p*=.487; study 3: *t*(49)=−0.234, two-sided *p*=.839).

Participants’ subjective reports of the perceived percentages of WIN and noWIN outcomes did not differ from 50% in studies 1 and 2 (study 1: LC: *M*=51.60%, *SD*=12.36, *t*(39)=0.818, two-sided *p*=.418, MC: *M*=48.23%, *SD*=6.60, *t*(39)=−1.70, two-sided p=.097, HC: *M*=50.70%, *SD*=14.78, *t*(39)=.300, two-sided *p*=.766; study 2: MC: *M*=48.13%, *SD*=6.37, *t*(37)=−1.808, two-sided *p*=.079, HC: *M*=52.07%, *SD*=10.29, *t*(37)=1.216, two-sided *p*=.232), nor were there any significant differences between conditions (study 1: *F*(2,78)=.982, two-sided *p*=.379; study 2: *t*(37)=−1.936, two-sided *p*=.061). Surprisingly, in study 3 subjects reported having received significantly less than 50% WIN outcomes in MC (*M*=46.66%, *SD*=11.75%, *t*(49)=−2.011, two-sided *p*=.049, *d*=−0.28, 90% *CI*=[−.57, −.0002]), and significantly more than 50% WIN outcomes in HC (*M*=54.51%, *SD*=12.07%, *t*(49)=2.640, *p*=.011, two-sided, *d*=0.37, 90% CI=[.09, .66]), contradicting the objective outcome rates. We therefore controlled the results of study 3 for differences between conditions regarding subjective outcome rates.

#### Analysis of affective Dynamics in Studies 1, 2, and 3

For each participant, we calculated the mean ratings of pride or happiness (study 3: only pride), separately for each combination of condition (LC (only study 1), MC, HC) and outcome valence (WIN, noWIN), for all valid trials (i.e. excluding those in which the participant either clicked on the background or did not respond fast enough). The resulting variables were then taken as dependent variables in repeated measures analyses of variance (rmANOVA), using the factors condition, outcome valence, and affect (factor affect only in studies 1 & 2). Paired comparisons were then used to disentangle significant effects of interest.

#### Correlation of Control Beliefs and Pride Response across Studies

In order to perform a more direct test regarding the relationship between internal control beliefs and pride, we computed the pride response from the data of each study ([HC:WIN–HC:noWIN] – [MC:WIN–MC:noWIN]). Hence, a positive pride response indicated that participants reported greater outcome-dependent differences in pride under HC than under MC. Additionally, after transforming the control ratings from each study to values between 0 and 100 across HC and MC, we computed the difference between control ratings for the different levels of control. For this analysis, control ratings and pride ratings from study 1 were averaged across LC and MC. Non-parametric correlation analysis (Spearman’s rho) was used to test the relationship between the pride response and differences in control beliefs between HC and MC.

#### Analysis of Choice Data in Study 3

We predicted the probability for each subject to choose the HC option on a given trial by fitting a logistic model to the choice data. We calculated the ratio of the potential monetary gain for the HC option (e.g. 1.37 €) to the one for the MC option (e.g. 1.01 €) on each of the 96 trials and transformed this ratio to log-space (e.g. log(1.37/1.01)=.3049). Thus, positive values of the resulting predictor variable indicate a higher potential gain for HC than for MC on a given choice trial, negative values indicate a relatively higher potential gain for MC, and a value of zero indicate equality of gains for the two options.

Considering the general form of logistic models,

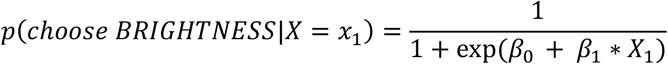

participants with parameters *β*_0_ and *β*_1_ both equaling zero would have a 50% probability of choosing HC on a given trial, regardless of the value of *X*_1_ (*X*_1_ being the log-transformed ratio of the monetary values offered for HC and MC). Assuming equal offers for HC and MC, negative values of *β*_0_ indicate that a participant has a preference for HC over MC, while positive values of *β*_0_ indicate a preference for MC over HC. With more negative values of *β*_1_, the probability of a participant choosing HC therefore increases more steeply, the higher the potential gain in the HC option, relative to MC.

Participants reported having received WIN outcomes more often in HC than in MC (see Supplement 1 - table S6), contradicting objective outcome rates. To control for differences in the subjective outcome rates, we again fit the logistic functions to each subjects’ choice data, now incorporating subjective outcome probabilities in the prediction. First, each monetary amount offered for either HC or MC on a given trial was multiplied with the subjectively rated percentage of WINs obtained in the respective condition (e.g. 1.23€ * 52% WINs in HC = 0.64€). The ratio of these expected values on each trial was then log-transformed and used to predict the subjects’ choices on each trial. The subjects’ probability to choose HC given equal expected values for HC and MC remained significantly above 50% (*MD*=65.98%, *IQR*=50.48, Wilcoxon signed-rank *W*=863, one-sided *p*=.015), i.e., even when controlling for differences in subjective winning histories for the two tasks, a preference for HC remained.

#### Analysis of FMRI Data (Study 2)

fMRI data were analyzed using SPM12^74^ in MATLAB R2015a. All functional volumes were slice-time corrected, spatially realigned, and normalized using the forward deformation fields as obtained using the unified segmentation approach by coregistration of the anatomical T1-image to the mean functional image of each individual participant^74^. Normalized images were resliced with a voxel size of 2*2*2 mm and smoothed with an 8-mm full-width-at-half-maximum isotropic Gaussian kernel. To remove drift, functional images were high-pass-filtered at 1/128 Hz.

Statistical analyses were performed in a two-level, mixed-effects approach. The first-level general linear model (GLM) for each participant included a total of 8 regressors defining the onsets and duration of the two task phases (MC, HC), the four feedback phases (MC:WIN, MC:noWIN, HC:WIN, HC:noWIN), and two regressors modelling the phases for pride and happiness ratings. In addition, the six realignment parameters from the preprocessing as well as their first derivatives were included as regressors of no interest to account for noise due to head movement.

For the analyses at the second level, we focused on the outcome phase of the task by implementing a within-subjects ANOVA including the individual first-level contrast images for MC:WIN, MC:noWIN, HC:WIN, and HC:noWIN. Using this approach we computed main effects of condition (HC>MC) and outcome valence (WIN>noWIN), as well as their interaction ((HC:WIN>HC:noWIN) > (MC:WIN>MC:noWIN)).

#### GLM Analysis including Pride and Happiness Ratings as parametric Modulators

We performed a second GLM analysis which included two predictors modelling the onsets and durations of the MC and the HC task on each trial, three different predictors for the task outcomes, and two predictors modelling the onsets and durations of the pride and happiness ratings. The first of the three outcome predictors modelled those task outcomes after which pride was rated. The second one modelled those task outcomes after which happiness was rated. For each of these predictors the respective affect ratings on each trial were included as parametric modulations to model variability in neural responses related to the subjective ratings of pride or happiness on each trial. The third outcome regressor modelled those trial outcomes that were not followed by any affect rating, and thus was not parametrically modulated. As before, the six realignment parameters as well as their first derivatives were included to account for noise due to head movement. The first-level contrast images for the parametric modulations of pride and happiness were then taken to a group-level within-subjects ANOVA in SPM12^74^.

#### Psychophysiological Interactions (PPI)

In addition to the standard GLM approach, we additionally performed psychophysiological interaction (PPI) analyses on the first level and aggregated the resulting contrast images for the PPI effect on the second level using one-sample t-tests in order to investigate changes in functional connectivity between brain regions identified in the within-participants ANOVA. Precisely, we investigated whether functional connectivity of regions activated more strongly for WIN than for noWIN would differ between outcomes of a HC as compared MC. For each individual participant, we defined 4-mm radius spherical ROIs, centered at the nearest local maximum for the contrast WIN>noWIN and located within 6 mm from the group maximum for the same contrast, separately for the left and right VS, which is a central structure associated with the processing of rewarding outcomes^24, 76^. By computing the first eigenvariate for all voxels within each of the two individual spheres that showed a positive effect for WIN>noWIN, we extracted the time course of activations and constructed PPI terms using the contrast HC>MC^74^. One participant was excluded from the PPI analysis for right VS, because no voxels survived the predefined threshold for eigenvariate extraction. The PPI term, along with the activation time course from the (left or right) VS and a regressor modelling the effect of HC>MC was included in a new GLM for each participant that additionally modelled the two task phases (grids for HC or MC) and rating phases (pride and happiness) as well as the six realignment parameters from preprocessing and their first derivatives. First level contrast images of the (left or right) VS PPI analyses were aggregated for all participants in order to test for changes in functional connectivity on the group level using one-sample t-tests.

#### Rationale for Masking Procedures

We created a set of masks used for region-of-interest (ROI) analyses by performing automated meta-analyses using Neurosynth^52^ for the terms *self referential* and *value*, comprising activation foci of 166 and 470 studies, respectively. Using the SPM12 image calculator^74^, we created three mutually exclusive image masks, showing brain regions associated with a) the term *self referential*, but not *value* (SELF¬VALUE), b) with the term *value*, but not *self referential* (VALUE¬SELF), and c) the conjunction of both terms (SELF∩VALUE, covering regions where both meta-analyses showed overlapping effects). This rationale is in line with a recent meta-analysis showing substantial overlap between regions associated with processing of SV and self-referential processes^77^. These masks were then limited to include only clusters with 5 or more contiguous voxels. For a given contrast, we first tested whether there were significant activations within the regions covered by the masks of interest. Subsequently, we tested whether the same contrast showed significant activations on the whole-brain level in order to detect regions activated that were located outside of our a-priori masks.

#### Linking Internal Control-dependent Functional Connectivity to interindividual Variability in affective Responses

We hypothesized that individuals showing stronger functional connectivity between VS and regions associated with self-related processing when receiving HC outcomes than when receiving MC outcomes, should display larger pride responses. We obtained an estimate of individual functional connectivity strengths by computing the eigenvariate across participants from the three largest clusters inside the SELF¬VALUE mask showing significant PPI effects at *p*<.0005 (uncorrected at peak level). These three clusters were located in left AG (*k*=50 voxels), dmPFC (*k*=89) and precuneus (*k*=33; figure 6). Next, we correlated pride responses with estimates of functional connectivity in each of the three clusters, applying FDR correction for multiple comparisons on the resulting *p*-values (using the *p.adjust* function in R^75^). An equivalent analysis was performed for the happiness responses. We then compared these correlations between pride and happiness using the *cocor* package^78^ in R, again applying FDR-correction for multiple comparisons.

## Supporting information

Supplemental tables for behavioral analyses

Supplemental tables for fMRI analyses

## Acknowledgements

We thank Janine Baumann, Finn Lübber, Timo Schlesinger, and Johanna Schulz for their help with data collection.

## Author Contributions

D.S.S., F.M.P., L.M.P., and S.K. developed and designed the experiments. D.S.S. programmed the experiments. D.S.S., F.M.P., and S.K. conducted data collection. D.S.S. performed statistical analyses. D.S.S., F.M.P., L.M.P., and S.K. interpreted the results. D.S.S., F.M.P., L.M.P., and S.K. wrote the paper.

