## Supplemental tables for behavioral analyses for "The Pursuit of Pride: Outcomes achieved under Beliefs of Internal Control shape positive Affect and neural Dynamics in the vmPFC"

**Supplement 1.** Supplemental information and statistics for behavioral analyses.

**Table S1**. Repeated measures ANOVA of affect ratings (study 1)

| Within subjects effect | *SSs* | *df* | *Mean Square* | *F* | *p* | $\eta_p^2$ |
|---|---|---|---|---|---|---|
| condition | 2018.4 | 1.31 | 1545.60 | 28.84[a] | <.001 | .43 |
| error | 2729.2 | 50.93 | 53.59 | | | |
| outcome valence | 79377.8 | 1.00 | 79377.77 | 60.92 | <.001 | .61 |
| error | 50819.7 | 39.00 | 1303.07 | | | |
| affect | 9904.4 | 1.00 | 9904.39 | 17.45 | <.001 | .31 |
| error | 22142.1 | 39.00 | 567.74 | | | |
| condition * outcome valence | 4760.4 | 1.37 | 3466.75 | 48.51[a] | <.001 | .55 |
| error | 3826.8 | 53.55 | 71.46 | | | |
| condition * affect | 389.4 | 1.31 | 297.16 | 12.87[a] | <.001 | .25 |
| error | 1180.4 | 51.11 | 23.10 | | | |
| outcome * affect | 9825.2 | 1.00 | 9825.19 | 70.10 | <.001 | .64 |
| error | 5466.4 | 39.00 | 140.16 | | | |
| condition * outcome valence * affect | 548.5 | 1.29 | 426.98 | 11.44[a] | <.001 | .23 |
| error | 1869.5 | 50.10 | 37.32 | | | |

**Note**. Type III sum of squares; SSs=Sum of Squares; Results are Greenhouse-Geisser corrected; [a] Mauchly's test indicates deviation from sphericity assumption.

**Table S2**. Repeated measures ANOVA of affect ratings (study 2)

| Within subjects effect | *SSs* | *df* | *Mean Square* | *F* | *p* | $\eta_p^2$ |
|---|---|---|---|---|---|---|
| condition | 328.1 | 1 | 328.06 | 8.38 | .006 | .18 |
| error | 1487.8 | 38 | 39.15 | | | |
| outcome valence | 132177.8 | 1 | 132177.83 | 174.83 | <.001 | .82 |
| error | 28728.8 | 38 | 756.02 | | | |
| affect | 998.3 | 1 | 998.26 | 12.69 | .001 | .25 |
| error | 2989.2 | 38 | 78.66 | | | |
| condition * outcome valence | 2517.4 | 1 | 2517.41 | 40.23 | <.001 | .51 |
| error | 2377.7 | 38 | 62.57 | | | |
| condition * affect | 119.7 | 1 | 119.75 | 3.24 | .080 | .08 |
| error | 1404.7 | 38 | 36.97 | | | |
| outcome * affect | 2009.8 | 1 | 2009.83 | 31.52 | <.001 | .45 |
| error | 2423.1 | 38 | 63.76 | | | |
| condition * outcome valence * affect | 249.3 | 1 | 249.26 | 9.08 | .005 | .19 |
| error | 1043.6 | 38 | 27.46 | | | |

**Note**. Type III sum of squares; SSs=Sum of Squares.

**Table S3**. Repeated measures ANOVA of pride ratings (study 3)

| Within subjects effect | *SSs* | *df* | *Mean Square* | *F* | *p* | $\eta_p^2$ |
|---|---|---|---|---|---|---|
| condition | 1917 | 1 | 1916.95 | 18.33 | <.001 | .27 |
| error | 5125 | 49 | 104.59 | | | |
| outcome valence | 30745 | 1 | 30744.92 | 136.19 | <.001 | .74 |
| error | 11062 | 49 | 225.76 | | | |
| condition * outcome valence | 3171 | 1 | 3171.08 | 35.98 | <.001 | .42 |
| error | 4319 | 49 | 88.15 | | | |

**Note**. Type III sum of squares; SSs=Sum of Squares.

**Table S4.** Reaction times and number of valid trials

| | | Median reaction times (ms) | | Number of valid trials | | Number of trials planned |
|---|---|---|---|---|---|---|
| | condition | *M* | *SD* | *M* | *SD* | |
| Study 1 | LC | 751 | 89 | 29.10 | 1.11 | 30 |
| | MC | 778 | 101 | 29.05 | 0.93 | 30 |
| | HC | 820 | 90 | 28.56 | 1.19 | 30 |
| Study 2 | MC | 1428 | 380 | 38.80 | 1.30 | 40 |
| | HC | 2041 | 400 | 38.13 | 1.99 | 40 |
| Study 3 | MC | 1223 | 325 | 30.76 | 1.24 | 32 |
| | HC | 1218 | 322 | 31.54 | 0.73 | 32 |

**Note.** ms = milliseconds; *M* = mean; *SD* = standard deviation; LC = low control; MC = medium control; HC = high control.

**Table S5.** Items regarding subjects' experience of the experimental task (study 3)

| Variable | Item |
|---|---|
| subjectively received %WIN | How often do you think that you guessed correctly / identified the brightest square in the different tasks? (in percent) |
| difficulty (inv.) | How easy were the different tasks for you? |
| effort | How much effort did you put into the different tasks? |
| fun | How much fun did you have doing the different tasks? |
| exciting | How exciting / thrilling do you think the different tasks were? |
| exhausting | How exhausting were the different tasks? |

**Table S6.** Rank correlation of pride response and choice preference (study 3)

| | | | statistics of control variables | | | | | |
|---|---|---|---|---|---|---|---|---|
| | | | HC | | MC | | | |
| | *rho* | $p^a$ | *M (SD)* | | *M (SD)* | | *t* | $p^b$ |
| Correlation of pride response with p(choose HC\|HC=MC) | .47 | <.001 | | | | | | |
| controlling for | | | | | | | | |
| subjectively received %WIN | .47 | <.001 | 54.51 | (12.07) | 46.66 | (11.75) | 5.34 | <.001 |
| difficulty | .49 | <.001 | 55.16 | (13.83) | 38.90 | (26.15) | 3.93 | <.001 |
| effort | .44 | <.001 | 81.26 | (12.50) | 45.05 | (28.03) | 9.02 | <.001 |
| exhausting | .42 | .001 | 62.15 | (17.51) | 21.65 | (24.18) | 10.01 | <.001 |
| fun | .41 | .002 | 62.58 | (17.92) | 36.83 | (18.96) | 8.12 | <.001 |
| exciting | .51 | <.001 | 58.13 | (23.14) | 30.45 | (22.82) | 7.07 | <.001 |

**Note.** *rho* = (partial) correlation coefficients computed using rank transformed variables; HC = high control; MC = medium control; *M* = mean; *SD* = standard deviation; a = one-sided; b = two-sided.
