## Supplemental tables for fMRI analyses for "The Pursuit of Pride: Outcomes achieved under Beliefs of Internal Control shape positive Affect and neural Dynamics in the vmPFC"

**Supplement 2.** Coordinates and full statistics for fMRI analyses.

**Table S7.** WIN>noWIN small-volume-corrected inside VALUE¬SELF, critical T = 3.881

| set | | cluster | | | | peak | | | | | MNI coordinates (mm) | | |
|---|---|---|---|---|---|---|---|---|---|---|---|---|---|
| *p* | *c* | *p* (FWE) | *p* (FDR) | *k* | *p* (unc) | *p* (FWE) | *p* (FDR) | *T* | *Z* | *p* (unc) | x | y | z |
| <.001 | 11 | <.001 | .002 | 235 | <.001 | <.001 | <.001 | 8.18 | 7.24 | <.001 | -14 | 8 | -10 |
| | | .002 | .108 | 61 | .039 | <.001 | <.001 | 8.14 | 7.21 | <.001 | -2 | 58 | 0 |
| | | | | | | <.001 | <.001 | 7.18 | 6.51 | <.001 | 2 | 56 | 0 |
| | | | | | | <.001 | .003 | 5.58 | 5.24 | <.001 | 4 | 50 | -6 |
| | | <.001 | .002 | 233 | <.001 | <.001 | <.001 | 7.69 | 6.89 | <.001 | 16 | 4 | -12 |
| | | <.001 | .032 | 108 | .009 | <.001 | <.001 | 6.71 | 6.15 | <.001 | -2 | 44 | -6 |
| | | | | | | .001 | .016 | 5.11 | 4.84 | <.001 | -4 | 44 | -12 |
| | | | | | | .001 | .030 | 4.93 | 4.68 | <.001 | 2 | 46 | -12 |
| | | .041 | .811 | 1 | .811 | <.001 | .002 | 5.83 | 5.44 | <.001 | 0 | 44 | 0 |
| | | .015 | .613 | 14 | .297 | <.001 | .003 | 5.68 | 5.32 | <.001 | -2 | -32 | 38 |
| | | .017 | .613 | 12 | .334 | <.001 | .003 | 5.59 | 5.24 | <.001 | 4 | 48 | -2 |
| | | | | | | <.001 | .004 | 5.48 | 5.15 | <.001 | 2 | 46 | 2 |
| | | | | | | <.001 | .005 | 5.40 | 5.08 | <.001 | 0 | 44 | 6 |
| | | .033 | .811 | 3 | .647 | .012 | .269 | 4.31 | 4.14 | <.001 | -2 | 40 | 10 |
| | | .036 | .811 | 2 | .718 | .012 | .269 | 4.30 | 4.13 | <.001 | -10 | 52 | -2 |
| | | .041 | .811 | 1 | .811 | .019 | .396 | 4.17 | 4.02 | <.001 | -2 | 48 | 10 |
| | | .041 | .811 | 1 | .811 | .033 | .659 | 4.01 | 3.87 | <.001 | -2 | 48 | -14 |

**Table S8.** WIN>noWIN small-volume-corrected inside SELF ∩VALUE, critical T = 4.866

| set | | cluster | | | | peak | | | | | MNI coordinates (mm) | | |
|---|---|---|---|---|---|---|---|---|---|---|---|---|---|
| *p* | *c* | *p* (FWE) | *p* (FDR) | *k* | *p* (unc) | *p* (FWE) | *p* (FDR) | *T* | *Z* | *p* (unc) | x | y | z |
| <.001 | 3 | <.001 | .001 | 111 | <.001 | <.001 | <.001 | 7.60 | 6.82 | <.001 | -2 | 48 | -4 |
| | | | | | | <.001 | .021 | 5.86 | 5.47 | <.001 | -6 | 46 | -4 |
| | | <.001 | .592 | 2 | .592 | <.001 | .005 | 6.29 | 5.82 | <.001 | -6 | 54 | -4 |
| | | <.001 | .592 | 3 | .503 | <.001 | .034 | 5.69 | 5.33 | <.001 | 0 | -34 | 36 |

**Table S9.** WIN>noWIN wholebrain p<.05, FWE-corrected

| set | | cluster | | | | peak | | | | | MNI coordinates (mm) | | |
|---|---|---|---|---|---|---|---|---|---|---|---|---|---|
| *p* | *c* | *p* (FWE) | *p* (FDR) | *k* | *p* (unc) | *p* (FWE) | *p* (FDR) | *T* | *Z* | *p* (unc) | x | y | z |
| <.001 | 12 | <.001 | <.001 | 175 | <.001 | <.001 | <.001 | 8.18 | 7.24 | <.001 | -14 | 8 | -10 |
| | | <.001 | <.001 | 598 | <.001 | <.001 | <.001 | 8.14 | 7.21 | <.001 | -2 | 58 | 0 |
| | | <.001 | <.001 | 1425 | <.001 | <.001 | <.001 | 8.12 | 7.20 | <.001 | 12 | -88 | 2 |
| | | | | | | <.001 | <.001 | 7.23 | 6.54 | <.001 | 12 | -76 | -10 |
| | | | | | | <.001 | .002 | 6.77 | 6.20 | <.001 | 14 | -94 | 22 |
| | | <.001 | <.001 | 175 | <.001 | <.001 | <.001 | 7.69 | 6.89 | <.001 | 16 | 4 | -12 |
| | | <.001 | <.001 | 163 | <.001 | <.001 | .018 | 6.15 | 5.70 | <.001 | 0 | -34 | 30 |
| | | | | | | .002 | .068 | 5.68 | 5.32 | <.001 | -2 | -32 | 38 |
| | | .005 | .127 | 19 | .095 | .001 | .039 | 5.90 | 5.50 | <.001 | -50 | -56 | -20 |
| | | <.001 | .005 | 78 | .002 | .001 | .039 | 5.87 | 5.47 | <.001 | 0 | -10 | 10 |
| | | <.001 | .015 | 54 | .009 | .002 | .062 | 5.72 | 5.35 | <.001 | 0 | -76 | 50 |
| | | .014 | .321 | 8 | .267 | .006 | .135 | 5.44 | 5.12 | <.001 | 28 | -40 | -48 |
| | | .002 | .062 | 30 | .041 | .010 | .217 | 5.30 | 5.00 | <.001 | 2 | -72 | 34 |
| | | .019 | .416 | 5 | .381 | .020 | .411 | 5.13 | 4.85 | <.001 | -34 | -78 | -22 |
| | | .025 | .503 | 3 | .503 | .030 | .617 | 5.01 | 4.75 | <.001 | -2 | -98 | 6 |

**Table S10.** HC>MC small-volume-corrected inside SELF¬VALUE, critical T = 4.027

| set | | cluster | | | | peak | | | | | MNI coordinates (mm) | | |
|---|---|---|---|---|---|---|---|---|---|---|---|---|---|
| *p* | *c* | *p* (FWE) | *p* (FDR) | *k* | *p* (unc) | *p* (FWE) | *p* (FDR) | *T* | *Z* | *p* (unc) | x | y | z |
| <.001 | 12 | <.001 | .004 | 208 | <.001 | <.001 | <.001 | 6.61 | 6.07 | <.001 | -8 | -58 | 20 |
| | | | | | | <.001 | <.001 | 6.55 | 6.02 | <.001 | -12 | -56 | 14 |
| | | | | | | <.001 | <.001 | 6.31 | 5.83 | <.001 | -6 | -56 | 14 |
| | | | | | | <.001 | <.001 | 6.30 | 5.82 | <.001 | -6 | -60 | 24 |
| | | .007 | .527 | 27 | .132 | <.001 | <.001 | 6.34 | 5.85 | <.001 | -8 | 50 | -12 |
| | | <.001 | .036 | 108 | .006 | <.001 | <.001 | 6.27 | 5.80 | <.001 | -2 | 38 | 4 |
| | | .014 | .649 | 14 | .270 | <.001 | .005 | 5.60 | 5.25 | <.001 | -6 | -50 | 6 |
| | | .040 | .798 | 1 | .798 | .001 | .049 | 5.01 | 4.75 | <.001 | -10 | 46 | -10 |
| | | .011 | .649 | 17 | .226 | .002 | .073 | 4.88 | 4.64 | <.001 | -48 | -70 | 26 |
| | | .035 | .798 | 2 | .699 | .003 | .098 | 4.78 | 4.56 | <.001 | -4 | 46 | -14 |
| | | .035 | .798 | 2 | .699 | .012 | .333 | 4.43 | 4.25 | <.001 | 0 | 48 | -14 |
| | | .032 | .798 | 3 | .626 | .013 | .333 | 4.41 | 4.23 | <.001 | 8 | 54 | 24 |
| | | .035 | .798 | 2 | .699 | .018 | .442 | 4.31 | 4.14 | <.001 | -8 | 50 | 16 |
| | | .040 | .798 | 1 | .798 | .026 | .574 | 4.22 | 4.06 | <.001 | 4 | 46 | -14 |
| | | .026 | .798 | 5 | .518 | .037 | .747 | 4.12 | 3.97 | <.001 | 6 | 50 | -14 |

**Table S11.** HC>MC small-volume-corrected inside SELF∩VALUE, critical T = 3.182

| set | | cluster | | | | peak | | | | | MNI coordinates (mm) | | |
|---|---|---|---|---|---|---|---|---|---|---|---|---|---|
| *p* | *c* | *p* (FWE) | *p* (FDR) | *k* | *p* (unc) | *p* (FWE) | *p* (FDR) | *T* | *Z* | *p* (unc) | x | y | z |
| .050 | 1 | .001 | .016 | 162 | .016 | <.001 | <.001 | 6.13 | 5.69 | <.001 | -8 | 48 | -12 |
| | | | | | | <.001 | <.001 | 5.81 | 5.42 | <.001 | -10 | 50 | -8 |
| | | | | | | <.001 | .020 | 4.62 | 4.41 | <.001 | 0 | 42 | 4 |
| | | | | | | <.001 | .020 | 4.60 | 4.40 | <.001 | 0 | 40 | 0 |
| | | | | | | .002 | .066 | 4.22 | 4.06 | <.001 | -8 | 44 | 0 |
| | | | | | | .004 | .118 | 4.00 | 3.86 | <.001 | 8 | 50 | -8 |
| | | | | | | .007 | .167 | 3.85 | 3.72 | <.001 | -6 | 54 | -4 |
| | | | | | | .010 | .213 | 3.73 | 3.62 | <.001 | -8 | 50 | 0 |
| | | | | | | .039 | .785 | 3.27 | 3.19 | .001 | -14 | 48 | -4 |

**Table S12.** HC>MC wholebrain p<.05, FWE-corrected

| set | | cluster | | | | peak | | | | | MNI coordinates (mm) | | |
|---|---|---|---|---|---|---|---|---|---|---|---|---|---|
| *p* | *c* | *p* (FWE) | *p* (FDR) | *k* | *p* (unc) | *p* (FWE) | *p* (FDR) | *T* | *Z* | *p* (unc) | x | y | z |
| <.001 | 13 | <.001 | <.001 | 392 | <.001 | <.001 | <.001 | 7.54 | 6.78 | <.001 | 2 | -70 | -34 |
| | | | | | | <.001 | .024 | 6.08 | 5.65 | <.001 | 12 | -68 | -18 |
| | | | | | | .004 | .159 | 5.56 | 5.22 | <.001 | 0 | -68 | -16 |
| | | <.001 | <.001 | 499 | <.001 | <.001 | .002 | 6.93 | 6.32 | <.001 | -8 | -56 | 14 |
| | | | | | | .017 | .518 | 5.17 | 4.89 | <.001 | -6 | -48 | 24 |
| | | <.001 | .018 | 59 | .007 | <.001 | .012 | 6.34 | 5.85 | <.001 | -8 | 50 | -12 |
| | | <.001 | .003 | 107 | .001 | <.001 | .012 | 6.33 | 5.85 | <.001 | 10 | -52 | 14 |
| | | <.001 | .004 | 93 | .001 | <.001 | .013 | 6.27 | 5.80 | <.001 | -2 | 38 | 4 |
| | | .006 | .229 | 16 | .123 | .017 | .518 | 5.16 | 4.88 | <.001 | -22 | -48 | -14 |
| | | .006 | .229 | 16 | .123 | .022 | .619 | 5.09 | 4.82 | <.001 | 58 | -28 | 26 |
| | | .022 | .629 | 4 | .435 | .025 | .619 | 5.06 | 4.80 | <.001 | 12 | -64 | -44 |
| | | .022 | .629 | 4 | .435 | .025 | .619 | 5.06 | 4.79 | <.001 | 56 | -52 | 6 |
| | | .025 | .654 | 3 | .503 | .033 | .780 | 4.98 | 4.73 | <.001 | 24 | -48 | -24 |
| | | .036 | .719 | 1 | .719 | .042 | .934 | 4.91 | 4.67 | <.001 | 16 | -60 | -48 |
| | | .036 | .719 | 1 | .719 | .047 | .949 | 4.88 | 4.64 | <.001 | -48 | -70 | 26 |
| | | .036 | .719 | 1 | .719 | .048 | .949 | 4.88 | 4.64 | <.001 | -30 | -34 | -18 |

**Table S13.** 3-way conjunction small-volume-corrected inside SELF ∩VALUE, critical T = 3.182

| set | | cluster | | | | peak | | | | | MNI coordinates (mm) | | |
|---|---|---|---|---|---|---|---|---|---|---|---|---|---|
| *p* | *c* | *p* (FWE) | *p* (FDR) | *k* | *p* (unc) | *p* (FWE) | *p* (FDR) | *T* | *Z* | *p* (unc) | x | y | z |
| .050 | 1 | .025 | .502 | 11 | .502 | .023 | .456 | 3.45 | 3.36 | <.001 | -4 | 48 | -10 |

**Note.** Effects listed are for the contrast [HC:WIN>HC:noWIN] ∩[HC:WIN>MC:WIN]∩[HC:WIN>MC:noWIN]

**Table S14.** parametric modulation HAPPY small-volume-corrected inside VALUE¬SELF, critical T = 4.237

| set | | cluster | | | | peak | | | | | MNI coordinates (mm) | | |
|---|---|---|---|---|---|---|---|---|---|---|---|---|
| *p* | *c* | *p* (FWE) | *p* (FDR) | *k* | *p* (unc) | *p* (FWE) | *p* (FDR) | *T* | *Z* | *p* (unc) | x | y | z |
| .050 | 1 | .028 | .556 | 3 | .556 | .022 | .434 | 4.55 | 4.04 | <.001 | 16 | 2 | -12 |

**Table S15.** parametric modulation HAPPY small-volume-corrected inside SELF ∩VALUE, critical T = 3.397

| set | | cluster | | | | peak | | | | | MNI coordinates (mm) | | |
|---|---|---|---|---|---|---|---|---|---|---|---|---|
| *p* | *c* | *p* (FWE) | *p* (FDR) | *k* | *p* (unc) | *p* (FWE) | *p* (FDR) | *T* | *Z* | *p* (unc) | x | y | z |
| .001 | 2 | .019 | .738 | 13 | .369 | .002 | .083 | 4.60 | 4.08 | <.001 | -2 | 48 | -4 |
| | | .042 | .835 | 1 | .835 | .021 | .419 | 3.74 | 3.43 | <.001 | -6 | 46 | -4 |

**Table S16.** parametric modulation HAPPY wholebrain p <.0001, uncorrected

| set | | cluster | | | | peak | | | | | MNI coordinates (mm) | | |
|---|---|---|---|---|---|---|---|---|---|---|---|---|
| *p* | *c* | *p* (FWE) | *p* (FDR) | *k* | *p* (unc) | *p* (FWE) | *p* (FDR) | *T* | *Z* | *p* (unc) | x | y | z |
| <.001 | 9 | .009 | .041 | 91 | .005 | .028 | .128 | 5.81 | 4.89 | <.001 | 8 | -88 | 6 |
| | | .047 | .105 | 53 | .023 | .177 | .410 | 5.08 | 4.41 | <.001 | -4 | 60 | 10 |
| | | | | | | .727 | .680 | 4.31 | 3.87 | <.001 | -14 | 64 | 10 |
| | | .201 | .327 | 24 | .109 | .255 | .454 | 4.92 | 4.30 | <.001 | 6 | -76 | -10 |
| | | .450 | .517 | 10 | .290 | .351 | .454 | 4.77 | 4.19 | <.001 | 20 | -88 | -14 |
| | | .335 | .445 | 15 | .198 | .470 | .573 | 4.61 | 4.08 | <.001 | -4 | 48 | -2 |
| | | .508 | .517 | 8 | .344 | .523 | .584 | 4.55 | 4.04 | <.001 | 16 | 2 | -12 |
| | | .794 | .766 | 1 | .766 | .696 | .680 | 4.35 | 3.89 | <.001 | 20 | -88 | 34 |
| | | .651 | .575 | 4 | .511 | .703 | .680 | 4.34 | 3.89 | <.001 | 14 | -98 | 8 |
| | | .651 | .575 | 4 | .511 | .754 | .680 | 4.28 | 3.84 | <.001 | 26 | -90 | 22 |

**Table S17.** parametric modulation PROUD small-volume-corrected inside VALUE¬SELF, critical T = 4.175

| set | | cluster | | | | peak | | | | | MNI coordinates (mm) | | |
|---|---|---|---|---|---|---|---|---|---|---|---|---|---|
| *p* | *c* | *p* (FWE) | *p* (FDR) | *k* | *p* (unc) | *p* (FWE) | *p* (FDR) | *T* | *Z* | *p* (unc) | x | y | z |
| <.001 | 7 | .009 | .801 | 22 | .175 | <.001 | .031 | 6.15 | 5.09 | <.001 | -2 | 56 | 0 |
| | | .023 | .801 | 7 | .444 | .002 | .159 | 5.34 | 4.58 | <.001 | 20 | 6 | -14 |
| | | .012 | .801 | 16 | .245 | .008 | .306 | 4.86 | 4.26 | <.001 | -2 | 44 | -6 |
| | | .040 | .801 | 1 | .801 | .031 | .953 | 4.35 | 3.90 | <.001 | -12 | 2 | -10 |
| | | .040 | .801 | 1 | .801 | .041 | .953 | 4.25 | 3.82 | <.001 | -16 | 10 | -8 |
| | | .035 | .801 | 2 | .703 | .043 | .953 | 4.23 | 3.81 | <.001 | 18 | 14 | -4 |
| | | .040 | .801 | 1 | .801 | .048 | .953 | 4.19 | 3.78 | <.001 | -20 | 10 | -6 |

**Table S18.** parametric modulation PROUD small-volume-corrected inside SELF¬VALUE critical T = 4.353

| set | | cluster | | | | peak | | | | | MNI coordinates (mm) | | |
|---|---|---|---|---|---|---|---|---|---|---|---|---|---|
| *p* | *c* | *p* (FWE) | *p* (FDR) | *k* | *p* (unc) | *p* (FWE) | *p* (FDR) | *T* | *Z* | *p* (unc) | x | y | z |
| .050 | 1 | <.001 | .003 | 122 | .003 | .001 | .020 | 5.96 | 4.98 | <.001 | -2 | 56 | 2 |
| | | | | | | .004 | .074 | 5.27 | 4.54 | <.001 | -2 | 62 | 10 |

**Table S19.** parametric modulation PROUD small-volume-corrected inside SELF ∩VALUE, critical T = 3.343

| set | | cluster | | | | peak | | | | | MNI coordinates (mm) | | |
|---|---|---|---|---|---|---|---|---|---|---|---|---|---|
| *p* | *c* | *p* (FWE) | *p* (FDR) | *k* | *p* (unc) | *p* (FWE) | *p* (FDR) | *T* | *Z* | *p* (unc) | x | y | z |
| <.001 | 3 | .003 | .163 | 96 | .054 | .001 | .036 | 4.83 | 4.24 | <.001 | -4 | 48 | -6 |
| | | | | | | .001 | .036 | 4.78 | 4.21 | <.001 | 0 | 42 | -6 |
| | | .043 | .870 | 2 | .802 | .002 | .045 | 4.58 | 4.06 | <.001 | -6 | 54 | -4 |
| | | .046 | .870 | 1 | .870 | .021 | .392 | 3.69 | 3.39 | <.001 | -8 | 50 | 0 |

**Table S20.** parametric modulation PROUD wholebrain p <.0001, uncorrected

| set | | cluster | | | | peak | | | | | MNI coordinates (mm) | | |
|---|---|---|---|---|---|---|---|---|---|---|---|---|---|
| *p* | *c* | *p* (FWE) | *p* (FDR) | *k* | *p* (unc) | *p* (FWE) | *p* (FDR) | *T* | *Z* | *p* (unc) | x | y | z |
| .001 | 7 | <.001 | .001 | 269 | <.001 | .008 | .061 | 6.15 | 5.09 | <.001 | -2 | 56 | 0 |
| | | | | | | .084 | .159 | 5.27 | 4.54 | <.001 | -2 | 62 | 10 |
| | | | | | | .221 | .301 | 4.86 | 4.26 | <.001 | -2 | 44 | -6 |
| | | .293 | .532 | 18 | .228 | .071 | .159 | 5.34 | 4.58 | <.001 | 20 | 6 | -14 |
| | | .014 | .032 | 102 | .009 | .084 | .159 | 5.27 | 4.54 | <.001 | 16 | -78 | -8 |
| | | | | | | .109 | .167 | 5.16 | 4.47 | <.001 | 10 | -88 | 0 |
| | | .371 | .532 | 13 | .304 | .424 | .498 | 4.54 | 4.03 | <.001 | -20 | 12 | -4 |
| | | .661 | .710 | 2 | .710 | .584 | .703 | 4.35 | 3.90 | <.001 | -12 | 2 | -10 |
| | | .661 | .710 | 2 | .710 | .668 | .762 | 4.25 | 3.82 | <.001 | -16 | 10 | -8 |
| | | .588 | .710 | 4 | .581 | .687 | .762 | 4.23 | 3.81 | <.001 | 18 | 14 | -4 |

**Table S21.** left nACC PPI small-volume-corrected inside SELF¬VALUE, critical T = 4.376

| set | | cluster | | | | peak | | | | | MNI coordinates (mm) | | |
|---|---|---|---|---|---|---|---|---|---|---|---|---|---|
| *p* | *c* | *p* (FWE) | *p* (FDR) | *k* | *p* (unc) | *p* (FWE) | *p* (FDR) | *T* | *Z* | *p* (unc) | x | y | z |
| .001 | 2 | .006 | .244 | 24 | .122 | .001 | .079 | 5.64 | 4.76 | <.001 | -46 | -70 | 38 |
| | | | | | | .003 | .079 | 5.42 | 4.62 | <.001 | -42 | -72 | 40 |
| | | .034 | .670 | 2 | .670 | .007 | .143 | 5.07 | 4.39 | <.001 | -38 | -72 | 40 |

**Table S22.** left nACC PPI small-volume-corrected inside SELF ∩VALUE critical T = 3.360

| set | | cluster | | | | peak | | | | | MNI coordinates (mm) | | |
|---|---|---|---|---|---|---|---|---|---|---|---|---|---|
| *p* | *c* | *p* (FWE) | *p* (FDR) | *k* | *p* (unc) | *p* (FWE) | *p* (FDR) | *T* | *Z* | *p* (unc) | x | y | z |
| <.001 | 4 | .033 | .863 | 5 | .649 | .010 | .863 | 4.00 | 3.62 | <.001 | -12 | 50 | 6 |
| | | .007 | .569 | 48 | .142 | .017 | .863 | 3.80 | 3.47 | <.001 | 6 | 48 | -14 |
| | | | | | | .026 | .863 | 3.62 | 3.33 | <.001 | 0 | 50 | -12 |
| | | | | | | .032 | .863 | 3.54 | 3.27 | .001 | -4 | 54 | -6 |
| | | .029 | .863 | 7 | .583 | .032 | .863 | 3.54 | 3.27 | .001 | -8 | 46 | 0 |
| | | | | | | .039 | .863 | 3.46 | 3.20 | .001 | -12 | 46 | 2 |
| | | | | | | .044 | .867 | 3.42 | 3.17 | .001 | -8 | 44 | -4 |
| | | .043 | .863 | 1 | .863 | .038 | .863 | 3.48 | 3.21 | .001 | -8 | 50 | 0 |

**Table S23.** left nACC PPI wholebrain p<.001, uncorrected, k >=50

| set | | cluster | | | | peak | | | | | MNI coordinates (mm) | | |
|---|---|---|---|---|---|---|---|---|---|---|---|---|---|
| *p* | *c* | *p* (FWE) | *p* (FDR) | *k* | *p* (unc) | *p* (FWE) | *p* (FDR) | *T* | *Z* | *p* (unc) | x | y | z |
| <.001 | 7 | .009 | .020 | 329 | .001 | .035 | .481 | 5.67 | 4.78 | <.001 | -44 | -70 | 40 |
| | | <.001 | <.001 | 3147 | <.001 | .171 | .612 | 5.02 | 4.36 | <.001 | 22 | 28 | 38 |
| | | | | | | .174 | .612 | 5.01 | 4.35 | <.001 | -8 | 46 | 50 |
| | | | | | | .249 | .612 | 4.85 | 4.24 | <.001 | -14 | 52 | 12 |
| | | .307 | .407 | 103 | .042 | .698 | .612 | 4.26 | 3.82 | <.001 | -44 | 24 | -14 |
| | | | | | | .765 | .612 | 4.18 | 3.76 | <.001 | -34 | 26 | -14 |
| | | .634 | .576 | 58 | .115 | .718 | .612 | 4.23 | 3.80 | <.001 | 46 | -64 | 42 |
| | | .625 | .576 | 59 | .112 | .775 | .612 | 4.16 | 3.75 | <.001 | -14 | 12 | 6 |
| | | .422 | .487 | 84 | .062 | .891 | .679 | 3.99 | 3.61 | <.001 | 0 | -56 | 16 |
| | | .054 | .083 | 207 | .006 | .914 | .690 | 3.94 | 3.58 | <.001 | -4 | -48 | 42 |
| | | | | | | .943 | .723 | 3.87 | 3.52 | <.001 | 4 | -56 | 36 |
| | | | | | | .983 | .740 | 3.71 | 3.40 | <.001 | 6 | -46 | 40 |

**Table S24.** right nACC PPI small-volume-corrected inside SELF¬VALUE, critical T = 4.373

| set | | cluster | | | | peak | | | | | MNI coordinates (mm) | | |
|---|---|---|---|---|---|---|---|---|---|---|---|---|---|
| *p* | *c* | *p* (FWE) | *p* (FDR) | *k* | *p* (unc) | *p* (FWE) | *p* (FDR) | *T* | *Z* | *p* (unc) | x | y | z |
| .001 | 2 | .026 | .766 | 4 | .512 | .028 | .821 | 4.59 | 4.07 | <.001 | -18 | 38 | 46 |
| | | .039 | .766 | 1 | .766 | .041 | .821 | 4.44 | 3.96 | <.001 | 0 | 36 | 32 |

**Table S25.** right nACC PPI wholebrain p<.001, uncorrected, k>=50

| set | | cluster | | | | peak | | | | | MNI coordinates (mm) | | |
|---|---|---|---|---|---|---|---|---|---|---|---|---|---|
| *p* | *c* | *p* (FWE) | *p* (FDR) | *k* | *p* (unc) | *p* (FWE) | *p* (FDR) | *T* | *Z* | *p* (unc) | x | y | z |
| .120 | 3 | <.001 | <.001 | 830 | <.001 | .078 | .205 | 5.35 | 4.59 | <.001 | -4 | 34 | 34 |
| | | | | | | .117 | .205 | 5.19 | 4.48 | <.001 | -22 | 38 | 46 |
| | | | | | | .887 | .910 | 4.01 | 3.64 | <.001 | -6 | 38 | 44 |
| | | .295 | .354 | 101 | .037 | .480 | .719 | 4.53 | 4.02 | <.001 | -38 | 14 | 54 |
| | | | | | | .998 | .910 | 3.54 | 3.27 | .001 | -32 | 14 | 44 |
| | | | | | | .998 | .910 | 3.54 | 3.27 | .001 | -44 | 16 | 46 |
| | | .668 | .744 | 53 | .118 | .810 | .910 | 4.13 | 3.73 | <.001 | -36 | 52 | 12 |
